# Microplastic pollution induces algae blooms in experimental ponds but bioplastics are less harmful

**DOI:** 10.1101/2025.10.24.683782

**Authors:** Scott G. Morton, Gabriel Vucelic-Frick, Jonathan R. Dickey, Bhausaheb S. Rajput, Cody J. Spiegel, Dahlia A. Loomis, Sara L. Jackrel, Michael D. Burkart, Jonathan B. Shurin

## Abstract

An ever-growing sea of plastic waste permeates even the most remote ecosystems; however, its ecological impact is unclear. Less persistent bioplastic alternatives are available but also have unknown environmental effects. We conducted a three-month experiment exposing plankton in experimental ponds to 10 concentrations of three different thermoplastic polyurethane microplastics, including two biodegradable bioplastics. Algal blooms with dense chlorophyll occurred consistently at high concentrations of the petroleum-derived thermoplastic polyurethane, but only occasionally with the two bioplastics. Herbivorous zooplankton density was strongly reduced by typical thermoplastic polyurethane and only weakly by bioplastics, therefore the effect on algae is at least partly due to reductions in top-down grazing pressure. Microbial communities exhibited compositional shifts in response to all three plastic types, with petroleum-derived plastic associated with the most pronounced differences across both prokaryotic and eukaryotic domains. Our results show that plastic pollution may contribute to the growing global problems of eutrophication, coastal hypoxia and harmful algae blooms, and that biodegradable plastics may have smaller environmental footprints.

## Introduction

Pervasive environmental pollution from plastic products and their eventual degradation into microplastics is a growing global concern ^1–3^. These pollutants have been detected in nearly all ecosystems, including remote and extreme environments such as deep sea trenches and arctic sea ice ^4,5^. Plastic debris undergoes physical, chemical and photodegradation into increasingly smaller particles that are transported by global processes such as ocean currents, hydrology, and trophic transfer ^6^. In freshwater systems, the ubiquitous presence of microplastics in the environment highlights their persistence and ability to spread far from their points of origin ^7–9^. Less than 1% of the 400 billion tons of plastic products produced annually are biodegradable ^10^ resulting in the continued accumulation of these pollutants.

Microplastics can integrate into the trophic networks of various aquatic ecosystems ^11–15^. These particles are capable of physically interacting with small organisms, such as zooplankton, and their increasing environmental concentrations may have cascading effects on consumers at both higher and lower trophic levels ^11,16–18^. These foreign particles can act as physical impediments and introduce chemical toxins, disrupting biological processes and impairing an organism’s fitness ^12,19^. Acute toxicity in aquatic organisms has been documented as a result of microplastic exposure ^20–23^. Some taxa, particularly microbes, can incorporate microplastics into their diets, and in certain cases, even convert them into usable, albeit recalcitrant, carbon sources ^24,25^. Beyond their direct effects on organisms, microplastic surfaces can also facilitate the proliferation of distinct microbial communities, including potential pathogens, which may further threaten aquatic taxa ^26– 29^. Yet, despite the ubiquity of microplastics in the environment, few studies have examined their broader ecological impacts, particularly in freshwater ecosystems, though exception can be found^19,30^.

One potential solution to the global plastic pollution problem is to devise new plastic formulations that are susceptible to microbial biodegradation and therefore less persistent in the environment ^31^. Bio-based plastics derived from algal lipids have been engineered to display similar material properties but can be fully biodegradable unlike most fossil-fuel based plastics ^32,33^. Although these materials may be less likely to accumulate over time, their environmental impacts at the ecosystem level are unknown.

Here we examine the effects of increasing concentrations of three types of microplastics on aquatic communities (i.e. zooplankton, algae, and bacteria) and ecosystem function (i.e chlorophyll-⍰, ecosystem respiration, and net primary production) in experimental ponds. We used one conventional petroleum-derived thermoplastic polyurethane (TPU), Elastollan, chosen for its commercial relevance; and two biodegradable TPU products, TPU 181 and TPU FC2.1, which represent emerging bio-based alternatives. The version of Elastollan that was tested is designed to be non-biodegradable as well as UV, hydrolysis, and microbial resistant ^34^. In contrast, products closely related to TPU 181 and TPU FC 2.1 have demonstrated 75% mass loss in composting conditions within 50 days ^35^, although degradation was not directly measured in our study. We used a gradient design with 10 unique concentrations of each of the three materials to test their impacts on aquatic organisms including bacteria, phytoplankton, protozoans and zooplankton, as well as effects on ecosystem trophic status and metabolism. Our experiment tests how the indirect effects of microplastics propagate across trophic levels including autotrophs, grazing consumers and microbes that recycle organic matter, and whether bioplastics have smaller environmental footprints than typical petroleum-based formulations.

## Results

### Chlorophyll-α

As a proxy for primary producer biomass, chlorophyll-α concentrations varied notably across plastic treatments over time (Fig. 1). The best fit generalized additive model (GAM) included plastic type as a fixed effect, with plastic type-specific smooth terms for time and concentration (Supplementary Table 1). Both bioplastics (TPU 181 and TPU FC2.1) yielded significantly lower chlorophyll-α (estimated effect sizes: = -0.567, -0.509, respectively) compared to the petroleum-derived plastic (reference level, Est. = 3.187) (Supplementary Table 2).

**Fig. 1:**
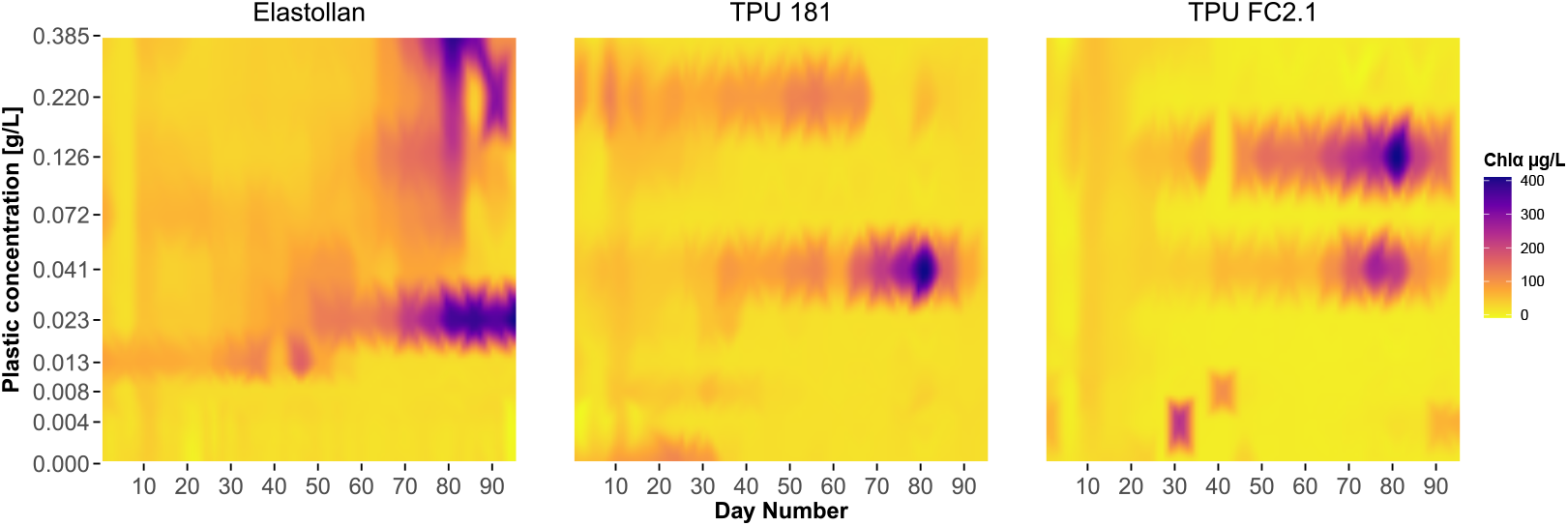
Chlorophyll-α concentrations across plastic types and concentrations over time. Heatmaps show interpolated chlorophyll-α concentrations for each plastic type across ten treatment concentrations (0.000–0.385 g/L) over the experiment duration. Interpolated values are based on the nearest chlorophyll-α readings, depicting a continuous gradient. Darker shades indicate higher chlorophyll-α concentrations.

The GAM model revealed significant non-linear effects of ‘plastic concentration’ and ‘sampling date’ on chlorophyll-α concentrations, varying distinctly across plastic types with the model explaining 56.9% (R^2^ = 0.547) of the deviance in chlorophyll-α concentrations (Supplementary Table 1; Supplementary Figure 1). Though algal blooms occurred with all plastic types at concentrations ≥ 0.041 g L⍰^1^, tanks treated with Elastollan above 0.013 g L⍰^1^ maintained consistently elevated chlorophyll-α throughout the second half of the experiment, an effect never seen at lower Elastollan concentrations (Fig 1).

### Net primary production and ecosystem respiration

Net primary production (NPP) exhibited distinct responses to plastic treatments (Supplementary Figure 2). The best-fit model included plastic type as a fixed effect and allowed for plastic-type-specific smooth terms for both time and concentration (Supplementary Table 1). Both bioplastics had significantly reduced NPP relative to the petroleum-derived plastic Elastollan (Est. = -0.200 for TPU 181, -0.102 for TPU FC2.1; Supplementary Table 2). Only Elastollan and TPU 181 yielded significant temporal trends in NPP, whereas effects of plastic concentration were only significant in ponds treated with Elastollan (p = 0.018), with no significant effect for either bioplastic (TPU 181: p = 0.082; TPU FC2.1: p = 0.177). The model explained 17.0% of the deviance (R^2^ = 0.147; Supplementary Table 1; Supplementary Figure 3).

Ecosystem respiration (ER) followed a similar pattern (Supplementary Figure 4), with reductions in both bioplastic treatments relative to Elastollan (Est. = –0.184 for TPU 181; –0.115 for TPU FC2.1; both p < 0.01). The chosen model included plastic-type-specific smooth terms for both time and plastic concentration, consistent with the structure selected for chlorophyll-α and NPP. Temporal trends were significant for TPU 181 and Elastollan, but not for TPU FC2.1, and a plastic concentration effect was observed only in Elastollan tanks (p = 0.029). The ER model explained 14.1% of the deviance (adjusted R^2^ = 0.113) (Supplementary Table 1; Supplementary Figure 5).

Together, these results suggest that exposure to plastic treatments altered both carbon fixation (NPP) and microbial respiration (ER), with the petroleum-derived plastic consistently associated with elevated metabolic activity relative to bioplastics.

### Zooplankton biomass and abundance

Elastollan-treated tanks exhibited significant zooplankton biomass deficits relative to both TPU 181 and TPU FC2.1 across a progressively narrowing concentration range over time (Fig. 2). On individual sampling dates, biomass variation was best explained by a model which included a fixed effect of plastic type and plastic-type-specific nonlinear effects of plastic concentration, indicating interactive effects between plastic type and concentration (Supplementary Table 3; Supplementary Figure 6). On Day 0 zooplankton biomass exhibited a statistically significant but visually weak nonlinear relationship with microplastic concentration (p = 0.026; Supplementary Table 4). As plastics had not yet been introduced to the system, this effect likely reflects random baseline variation. Supporting this interpretation, separate analyses found no significant differences among treatments in total biomass (p = 0.39), Shannon diversity (p = 0.647), or community composition (p = 0.928). Additionally, the model with a single smooth term for plastic concentration had the lowest AIC on Day 0 (Supplementary Table 3), further suggesting that observed variation was due to baseline noise rather than treatment effects. After the addition of plastic, the effects of plastic type and concentration were significant predictors of zooplankton biomass (Supplementary Table 4). Across sampling dates, Elastollan consistently reduced zooplankton biomass relative to both bioplastics, with the magnitude of this effect increasing at higher concentrations (Supplementary Table 5). On Day 36, model-predicted biomass in Elastollan-treated tanks was approximately 65% lower than TPU 181 and 69% lower than TPU FC2.1. These differences grew over time, reaching 71– 79% lower biomass by Day 56, and remaining 71–73% lower by Day 97. On Day 36, significant reductions in ln-biomass were observed at concentrations as low as 0.008 g□L^−1^, but by Day 97, only concentrations ≥□0.072 g□L^−1^ remained significant for TPU 181 and concentrations ≥□0.126 g□L^−1^ for TPU FC2.1. This shift toward higher concentration thresholds, despite consistently large biomass deficits, suggests that Elastollan’s suppressive effect on zooplankton biomass weakened as the experiment progressed (Supplementary Table 5).

**Fig. 2:**
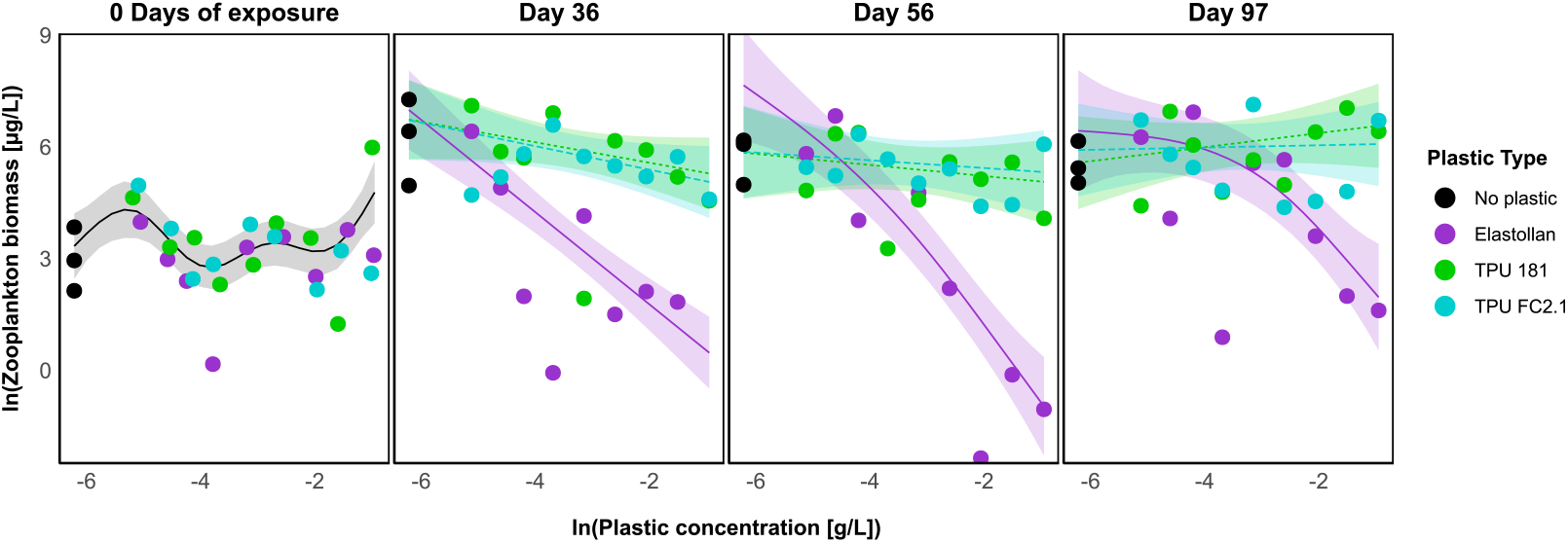
Zooplankton biomass across plastic concentrations and exposure times. Biomass of common zooplankton (µg/L) plotted against plastic concentration (g/L) for three plastic types before plastic application (left panel, 0 days of exposure) and at three subsequent time points (right panels). All lines represent the estimated biomass across concentrations, with shaded areas indicating 95% confidence intervals.

Abundance responses broadly mirrored biomass patterns, though effects were less pronounced and emerged later (Supplementary Figure 7, 8). As with biomass, reductions in abundance were concentration-dependent and varied by plastic type. At Day 0, no differences in abundance were detected across treatments (Supplementary Table 4), and Model 1 was selected to represent baseline variation. Following exposure, plastic type and concentration jointly explained significant variation in zooplankton abundance (Supplementary Table 4). Elastollan-treated tanks showed the strongest and most consistent reductions in abundance, with detectable effects from day 36 onward. These patterns were accompanied by shifts in the relative abundance of dominant taxa, suggesting community-level impacts (Supplementary Fig. 9).

### Microbial communities

We analyzed microbial community composition from 16S and 18S sequencing data to test the effects of plastic type and plastic concentration across multiple time points (Fig 3, Supplementary Fig 10-13). The PERMANOVA analyses for both 16S and 18S datasets identified significant main effects of plastic type, date, and chlorophyll-⍰ on community composition (Supplementary Table 6). The interaction of plastic concentration and plastic type was also significant. Plastic type explained a small but significant portion of the variation in both datasets (16S: R^2^ = 0.030, p = 0.004; 18S: R^2^ = 0.042, p = 0.003), while date consistently accounted for the largest amount of variation (16S: R^2^ = 0.157; 18S: R^2^ = 0.167; p < 0.001 for both) (Supplementary Table 7). Chlorophyll-⍰ was a significant covariate in both datasets, reflecting its role in shaping microbial community composition across sampling periods. Plastic concentration was significant in the 16S dataset (p = 0.030) but not in the 18S dataset (p = 0.117), possibly suggesting bacterial taxa may respond more strongly to resources or surface availability associated with plastic concentration than eukaryotic taxa.

**Fig. 3:**
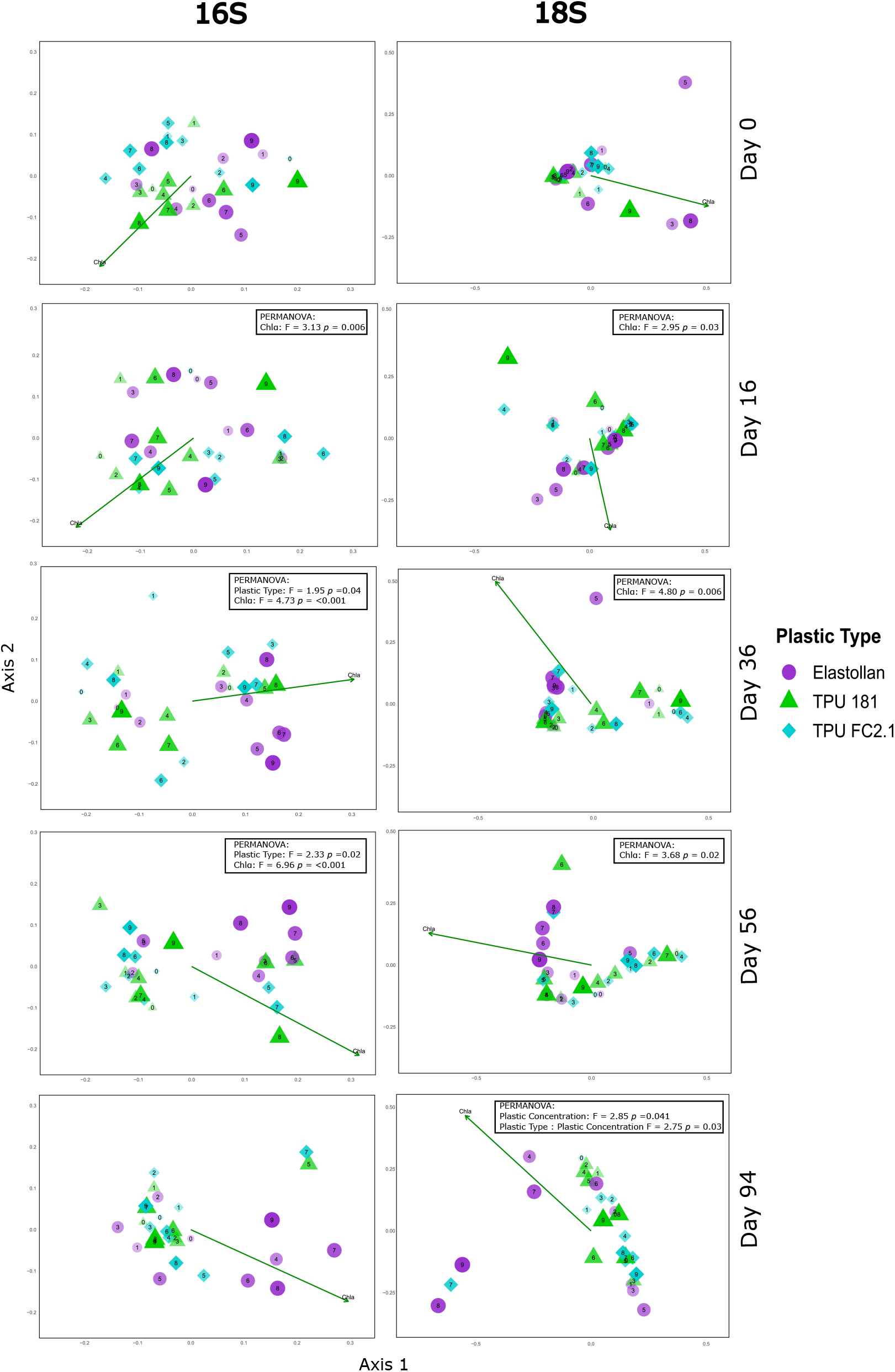
PCoA ordinations of 16S and 18S microbial communities. Principal coordinates analysis (PCoA) of microbial community composition based on 16S (left) and 18S (right) communities at 0 (pre-addition), 16, 36, 56 and 94 days of plastic exposure. Each point represents one tank: shape and color denote plastic type (Elastollan, TPU 181, TPU FC2.1), and the numeric label (0–9) indicates plastic concentration. Symbol size increases and transparency decreases with rising concentration. The green arrow shows the direction and strength of the chlorophyll-α vector in ordination space. Significant PERMANOVA results (p < 0.05) are overlaid to highlight treatment-driven shifts in community structure. The specific plastic concentrations (g/L) corresponding to labels 0–9 are as follows: 0 = 0.000, 1 = 0.004, 2 = 0.010, 3 = 0.013, 4 = 0.023, 5 = 0.041, 6 = 0.072, 7 = 0.126, 8 = 0.220, and 9 = 0.385.

Date-specific PERMANOVA analyses confirmed no significant differences in community composition across plastic treatments on day 0 for either dataset, indicating that subsequent shifts were treatment-induced rather than due to initial variation (Supplementary Table 7). For the 16S dataset, plastic type significantly affected community composition on days 36 (p = 0.037) and 56 (p = 0.022), while plastic concentration showed a marginal effect on day 56 (Supplementary Table 7). Chlorophyll-⍰ was a significant covariate on days 16, 36, and 56. In the 18S dataset, plastic type was marginally significant on day 56, while plastic concentration significantly influenced community composition on day 94 (p = 0.041), along with a significant interaction between plastic type and concentration (p = 0.032; Supplementary Table 7). Chlorophyll-⍰ remained a significant covariate on days 16, 36, and 56. Treatment-specific clustering and temporal shifts in community composition, as visualized in the PCoA, corroborated the PERMANOVA findings (Fig. 3).

To further investigate the taxa contributing to community shifts, we overlaid biplots of the ten most abundant families that significantly correlated with community composition onto PCoA plots for the 16S and 18S datasets on days 36, 56, and 94 (Supplementary Fig. 14, 15). These biplots revealed distinct responses among bacterial and eukaryotic taxa, with clustering patterns indicating that plastic type influenced microbial community structure across multiple domains of life. Additionally, we performed differential abundance analysis using DESeq2 on the 18S dataset for the final two sampling dates (days 56 and 94) when blooms were most apparent, to identify taxa driving shifts in community composition (Supplementary Fig. 16). On day 56, taxa enriched in Elastollan tanks included both phytoplankton and heterotrophic protists, such as members of the family Ochromonadaceae (golden algae), the order Ulotrichales (green algae), and the families Paramoebidae and Tetramitia (amoeboid protists) (Supplementary Fig. 16). In contrast, both bioplastic treatments (TPU 181 and TPU FC2.1) were associated with enrichment of Pleurostomatida (predatory ciliates), Maxillopoda and Branchiopoda (crustacean zooplankton), as well as Chilodonellidae and Sphaeropleales, which include some algal taxa. These enrichments occurred despite lower overall algal biomass, suggesting that bioplastic exposure may have favored communities shaped by top-down grazing pressure rather than algal proliferation.

By day 94, Elastollan ponds continued to show enrichment of phototrophic and mixotrophic taxa, including Chlorellales (green algae) and Paraphysomonadaceae (mixotrophic flagellates), consistent with sustained bloom conditions.These results indicate that Elastollan exposure influenced not only algal communities but also broader eukaryotic assemblages critical to microbial food webs. In contrast, bioplastic treatments were enriched in Chlamydomonadales, an early colonizer^36^, as well as Branchiopoda and Gregarinidae (parasitic protists), again suggesting a community structure potentially shaped by grazing rather than bloom dominance.

## Discussion

Our experiment suggests a functional link between two ongoing global trends in water pollution: eutrophication and microplastic contamination. Nutrient pollution has led to a proliferation of harmful algae blooms, deoxygenation, and impairment of freshwater resources ^37-39^, with implications for human health ^40^ and biodiversity ^41,42^. Eutrophication is commonly understood as a bottom-up phenomenon driven by excess nutrients; however, warming and loss of grazing zooplankton also contribute ^37–39 43^. The ecological impacts of microplastics are only beginning to be studied ^19,30,44^. Our experiment shows that high concentrations of fossil-fuel derived microplastics can stimulate algal blooms in experimental ponds, reduce abundance and biomass of grazing zooplankton and shift the composition of microbial communities. Bioplastics mitigated the first of these two impacts, with dampened effects on phytoplankton and zooplankton. Our results suggest that microplastics may facilitate cascading effects on the trophic structure of pelagic ecosystems by disrupting grazers control of algae.

Harmful algal blooms (HABs) are stochastic, unpredictable events typically emerging from nonlinear interactions, environmental drivers, and biological communities ^45–47^. In our experiment, dense algae concentrations always developed in tanks with high concentrations of Elastollan, but never in the three tanks with the lowest levels of microplastics. However, these blooms were relatively asynchronous, following differing trajectories, and their intensity was not clearly related to a dose-response relationship above the lower threshold. In each of the two bioplastics treatments, blooms developed in tanks treated with plastic concentrations = 0.041g/L, and additionally at 0.126g/L with TPU FC2.1, but not in any other tanks. Our results indicate that microplastics may tip the balance of conditions in favor of algal blooms but that stochasticity still plays a large role in when and where such blooms occur.

Here we show experimental ponds containing bioplastics resulting in greater zooplankton biomass compared to a petroleum-derived product. Elastollan, like many other commercial plastics, claims to have little to no acute negative impacts on freshwater organisms ^34^. Yet, in comparison to zooplankton populations exposed to either bioplastic alternative, Elastollan had a significant and consistent negative impact on grazer populations (Supplementary Figure 3, 4). Zooplankton biomass also declined with increasing concentrations of both TPU 181 and TPU FC2.1 on day 36, but by day 56, only TPU 181 continued to show a significant negative effect. By day 97, neither bioplastic had a detectable effect on zooplankton biomass, suggesting temporary or concentration-limited impacts. In contrast Elastollan’s negative effect remained statistically significant throughout the experiment, though the threshold concentration associated with suppression increased over time. Across the full 97-day period, zooplankton biomass in Elastollan treated ponds never recovered to levels observed in bioplastic treatments. This disruption in top-down control mediated by plastic pollution led to ecosystem scale effects as seen in ponds with rapid phytoplankton proliferation.

Our findings suggest microplastics play a role in shaping microbial community composition, with effects varying by plastic type and concentration. This supports previous studies showing that plastic pollution can alter nutrient cycling, microbial colonization, and ecosystem functioning beyond its physical presence ^19,27^. The significant effects of plastic type, concentration, and date on communities in both the 16S and 18S datasets suggest that plastics’ impacts vary among organismal groups, driving shifts in community composition over time. Importantly, microbial communities in no-plastic control tanks remained relatively stable across sampling periods, indicating that the observed shifts were not simply due to background temporal variation but were likely driven by plastic exposure (Supplementary Table 8). The observed association between chlorophyll-⍰ and both prokaryotes and eukaryotes underscores the strong coupling between microbial dynamics and algal blooms, suggesting that plastic-induced disruptions to trophic interactions - such as reduced grazer control of algae - cascade through biotic communities. Plastic-associated factors structured microbial communities beyond just algal taxa. The significant clustering of multiple bacterial and eukaryotic taxa in relation to plastic type and concentration indicates that Elastollan exposure altered community assembly processes, with potential implications for microbial succession and biofilm formation in the experimental ponds. These shifts in microbial composition suggest that petroleum-derived plastics like Elastollan not only promote algal proliferation but also restructure microbial food webs across multiple trophic levels. The consistent enrichment of golden algae (Ochromonadaceae) and green algae (Ulotrichales, Chlorellales) in Elastollan treatments indicates a plastic-specific selective effect on bloom-forming phytoplankton taxa. Simultaneously, the increased abundance of heterotrophic and mixotrophic eukaryotes, including Tetramitia and Paraphysomonadaceae highlights an expansion of microbial grazers and decomposers, which could be in response to enhanced algal biomass. Notably, Tetramitia, a free-living amoebae involved in bacterial predation were enriched in later sampling points, potentially responding to bacterial community changes. Among prokaryotes, the association of Acetobacteraceae, Comamonadaceae, and Caulobacteraceae with Elastollan treated ponds suggests plastic-altered carbon and nitrogen cycling pathways, as these taxa are known for their roles in organic compound degradation and biofilm formation^48-50^. The asynchronous effects of plastic type and concentration on microbial communities, observed at specific time points, highlight the temporal dynamics of ecosystem responses to plastic pollution across multiple domains of life. These results collectively illustrate that microplastics, particularly petroleum-derived plastics, may destabilize microbial community structure and function, amplifying the ecological impacts of eutrophication in freshwater systems.

The present study reveals that biodegradable plastics have smaller impacts on zooplankton communities than petroleum-derived plastics, a major regulating mechanism of harmful algal blooms ^51,52^. Additionally, the consistent appearance of algal blooms in experimental ponds containing the petroleum-derived plastic as compared to the infrequent presence in the tanks with biodegradable plastics reveals the potential environmental harm inflicted by plastic waste. The reduced biomass and abundance of zooplankton and frequent algal blooms at high plastic concentrations suggest that plastic pollution is a threat to the integrity of aquatic ecosystems, potentially contributing to eutrophication, oxygen depletion and harmful algal blooms in lakes and coastal zones. Transitioning to a biodegradable plastics economy is likely to mitigate the environmental impact of plastics in aquatic ecosystems.

## Materials and methods

Thirty experimental ponds were used to test the differential effects of three types of microplastics, one petroleum-derived thermoplastic polyurethane (TPU) an ester-based product (BASF Elastollan® #1175A10W), and two biodegradable TPUs (TPU 181, prepared from an algae source with ∼8.26% algae content, and TPU FC2.1; synthesized by B. S. Rajput [formulation in supplementary text] and Algenesis Materials Inc. [similar to ^53^] respectively), both made from compounds that can be derived from lipids produced by algae ^53^. Shore A hardness for each plastic were measured as follows: 75 (Elastollan)^54^, 88 (TPU FC2.1)^55^, and 97 (TPU 181; Supplementary methods). Each pond was a 400L elliptical tank (125cm x 65cm x 60cm; major axis x minor axis x depth) made of high-density polyethylene (HDPE) and placed outdoors under ambient temperature and light conditions (University of California, San Diego Biology Field Station, San Diego, CA). The tanks had been in use for several years prior to the experiment, and any leaching of plastic additives from the HDPE material was likely minimal. In a gradient design, each experimental pond contained increasing concentrations of one type of microplastic (by a factor of x1.75 g/L), across 10 concentrations ranging from 0-0.385 g/L. Ponds were placed in a grid formation and treatments were assigned randomly to reduce spatial bias. Tanks remained unstirred for the duration of the experiment, with minimal agitation occurring during routine sampling periods. To control diurnal variability, all distinct sampling efforts were conducted at approximately the same time of day (±2 hours), allowing for logistical constraints.

The range in microplastic concentration was determined based on plastic pollution observed globally ^56,57^. In the literature, microplastic concentrations are commonly reported in items/volume[area] ^58^. To relate these observed concentrations to our experimental treatments (expressed as weight per volume) we standardized the amount of plastic added by referencing each TPU as having a density of 1.2 g/cm^3 59^ and spherical particles for any size class. We used the following equations to convert from environmentally observed items/L to g/L.

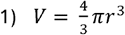

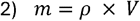

**Equation 1:** For a spherical particle, the volume (*V*) can be calculated as shown where (*r*) is the radius of the sphere.

**Equation 2:** The mass (*m*) of the particle is calculated from the density of the particle (*p*) multiplied by the volume (*V*).

To produce microplastics we combined plastic pellets with dry ice and continually blended them in a Nutribullet Magic Bullet blender (Nutribullet LLC, Los Angeles) with small holes in the lid to release CO_2_. These microplastic particles were then sieved into three distinct size classes ranging from 1000-711μm, 710-351μm, and <350 μm. Plastic particles in each size class were added to experimental ponds in a ratio of 58:25:17 percent, based on the total treatment weight. The particle size ratio added to each pond reflected the relative abundances of each size class available following processing and sieving of the materials. This ratio was standardized across all treatments to maintain consistency.

Each experimental pond was filled with municipal water (Nov 7, 2022), San Diego, CA and seeded with a concentrated mixture of live zooplankton, phytoplankton and microbes. Background environmental parameters were measured prior to the start of the experiment (Supplementary Table 9), with additional water quality metrics reported by the City of San Diego Public Utilities Department^60^. The community inacula was collected using vertical tows of a >63 μm mesh plankton net from Lake Miramar (Nov 11, 2022) and Lake Murray, San Diego, CA (Nov 22, 2022, and Jan 6, 2023) with 25 mL added on Nov 11 and Nov 22, and 35 mL on Jan 6. Inoculate volume reflected variation in the total plankton yield from each field sampling event. Multiple inoculations were performed to help establish robust zooplankton populations prior to microplastic additions. To support algal growth of phytoplankton from the inoculate and ensure a sufficient food source for zooplankton, nutrients were added to each tank prior to the start of the experiment. Tanks were enriched with 25 μg/L (Dec 26, 2022) and 100 μg/L (Dec 29, 2022) of nitrogen and phosphorus (10:1 molar ratio) with sodium nitrate (NaNO_3_) and sodium phosphate dibasic (Na_2_HPO_4_)respectively. Plastics were then added to each experimental pond (Mar 9, 2023) at specified treatment concentrations.

We sampled environmental parameters (temperature, DO, pH, conductivity, TDS, and salinity) using a YSI Quatro handheld multiparameter water sensor (YSI Inc., Yellow Spring, OH) calibrated against certified standards. Measurements of dissolved oxygen concentrations (% DO) were used to calculate net primary production (NPP) and ecosystem respiration (ER) at 8 time points (Starting on day 1, 6, 16, 26, 36, 56, 76, and 97). NPP was estimated as the change in % DO from dawn to dusk, averaged over three sequential daylight periods. ER was calculated as the change in % DO from dusk to the following dawn, averaged over two sequential nighttime periods. On sampling days 75–77, one daytime measurement (day 75) was missed. As a result, NPP was averaged over two daylight periods and ER was based on a single nighttime period for this time point. Data points where NPP < 0 or ER > 0, were considered probable sampling errors and excluded from further analysis. In-vivo chlorophyll-⍰ (a proxy for standing algal biomass) was measured every 5 days with a cuvette modular fluorometer/spectrophotometer (Turner Trilogy, Turner Designs, Sunnyvale California, USA) calibrated with a liquid dye standard (Turner Designs, Rhodamine WT Dye, 400 ppb, Sunnyvale California, USA). Zooplankton were sampled 4 times; 48 hours before plastics were added (March 7, 2023), on day 36 (April 23, 2023), on day 56 (May 3, 2023), and finally on day 97 (June 12, 2023). Zooplankton were collected by sampling 4L from each tank using a vertically integrated water sampler, which captured the entire water column, and preserved in 70% ethanol. Zooplankton were then identified to the lowest possible taxon using available published keys^61^ and photos were taken of a maximum of 30 individuals of each species and converted to biomass using length-mass regressions.

Microbial (16S and 18S) communities were sampled from each experimental pond five times over the course of the experiment; 48 hours prior to plastics addition (March 7, 2023), on day 16 (March 23, 2023), day 36 (April 23, 2023), day 56 (May 3, 2023), and on day 94 (June 9, 2023). Water samples from each pond were collected using a vertical integrated water sampler, in synergy with zooplankton samples, water was taken from both sides and the center of each pond, transferred and homogenized in sterile 5-gallon plastic buckets. Using a MasterFlex peristaltic pump (Masterflex L/S Portable Sampling Pump, Avantor, Radnor, Pennsylvania, USA), 2-3 liters were filtered through a 47mm, 0.22μm Millipore Express PLUS membrane filter (Merck KGaA, Darmstadt, Germany). Filters were immediately frozen in liquid nitrogen prior to storage at -80°C. For each sample, DNA was extracted from half of the filter following the QIAGEN DNeasy® PowerSoil® Pro Kit protocol (DNeasy PowerSoil Pro Kit Handbook, Qiagen, Hilden, Germany). For each sample, amplicon libraries targeting the V4-V5 region of the 16S rRNA gene and V9 region of the 18S rRNA gene were generated following the Earth Microbiome Project protocol at the Argonne National Lab’s Environmental Sample Preparation & Sequencing Facility. Library preparation was conducted using the 515-806R primer set for 16S and 1389F-1510R primer set for 18S, and amplicons were sequenced on a 2×151bp paired-end NextSeq 2000 run.

All sequence processing and quality control were conducted in QIIME2. Raw reads were demultiplexed, quality filtered, and denoised using the DADA2 plugin. For 16S amplicon sequences, paired-end reads were truncated to remove primers (forward = 138 bp, reverse = 130 bp) and filtered based on quality scores (maxEE forward = 2, reverse = 3). Reads were processed using a pseudo-pooling method and chimeric sequences were removed using the consensus method. For 18S rRNA amplicon sequences, paired-end reads were truncated to remove primers (forward = 140 bp, reverse = 138 bp) and filtered using quality scores (maxEE forward = 2, reverse = 2). Reads were processed using an independent pooling method and chimeric sequences were removed using the consensus method.

Amplicon sequence variants (ASVs) were inferred after merging, paired end reads using the *mergePairs* function in DADA2. Taxonomic classification of 16S sequences was performed against the SILVA 138 reference database using the naive Bayes classifier. 18S sequences were classified against PR2 trained on the V9 region, following the classifier training steps (including exact sequence matching and naïve Bayes classification). A phylogenetic tree was constructed by globally aligning ASVs using MAFFT v7 and inferring tree topology with FastTree (v2.1.4) under a GTR-CAT model.

The resulting 16S and 18S datasets were imported into R and processed using the *phyloseq* package ^62^ to construct a unified microbiome dataset. To ensure the removal of non-target sequences from the 16S dataset, a series of filtering steps were applied using the *subset_taxa* function in *phyloseq*. We first excluded ASVs lacking Kingdom-level assignments designated as unassigned, followed by the removal of sequences assigned to *Eukaryota*. Only ASVs classified as *Bacteria* or *Archaea* were retained for further analysis. To reduce non-target contaminants, we excluded sequences assigned to *Chloroplast* at the Order level and *Mitochondria* at the Family level. Finally, we filtered out poorly classified ASVs lacking reliable Phylum-level annotations.

A similar approach was applied to the 18S dataset. Taxonomic assignments were examined at multiple hierarchical levels to identify and remove bacterial, mitochondrial, and plasmid-derived sequences. Initially, sequences assigned to the Domain *Bacteria*, as well as unassigned sequences, were removed to focus the analysis on eukaryotic taxa. Next, sequences identified as mitochondrial or plasmid-derived DNA (categorized as *Eukaryota:mito* and *Eukaryota:plas*) were excluded. Further filtering involved removing unassigned taxa at additional hierarchical levels (Super Group, Super Kingdom, Kingdom, Phylum, Class, and Family) to reduce taxonomic ambiguity. Sequences classified as *Fungi* were excluded at the Kingdom level, as 18S rRNA gene markers are less reliable for fungal identification compared to more appropriate markers such as the Internal Transcribed Spacer (ITS) region, which was not used in this study. Three samples from day 0 were removed from the analysis due to their extremely low read depths. This stepwise approach ensured that downstream analyses focused on well-characterized eukaryotic taxa with ecological relevance to the research questions.

The final 16S dataset contained only bacterial and archaeal sequences, while the final 18S dataset consisted of eukaryotic taxa which were subsequently used for downstream analyses. To account for differences in sequencing depth across samples, we performed repeated rarefaction using the *phyloseq_mult_raref_avg* function from the *metagMisc* package ^63^. Each sample was randomly subsampled without replacement 100 times (i.e., rarefied), and ASV counts were averaged across iterations to reduce variation inherent to single rarefaction. Rarefaction was performed to a fixed depth based on the minimum sequencing depth across samples, 41,255 reads for 16S and 18,373 reads for 18S.

### Statistical Tests

We randomly assigned one unique control tank (0 µg/L microplastic) to each plastic type at the outset of the experiment to maintain a balanced factorial design. This approach ensured that each level of plastic type in the statistical models included a corresponding 0‐concentration reference, avoiding confounding between plastic type and concentration. While some natural variability among controls was observed (as is expected in outdoor systems), these differences were minor relative to treatment effects.

The effects of sampling date, plastic type, and plastic concentration on chlorophyll-⍰ concentrations, NPP, and ER were analyzed using a generalized additive model (GAM) implemented in the *mgcv* package in R ^64,65^. These data were manually log-transformed to improve distributional assumptions. We compared four models using Akaike Information Criterion (AIC). The simplest model (Model 1) included smooth terms for sampling date and raw plastic concentration, representing global effects across all plastic types. (Model 2), included plastic type as a parametric term, allowing for treatment-specific intercepts but assuming shared smoothers for sampling date and plastic concentration. (Model 3), allowed the smooth terms for sampling date and plastic concentration to vary by plastic type, providing treatment-specific smoothers. Finally, (Model 4) included treatment-specific smoothers for both sampling date and plastic concentration, with separate smooth functions fitted to each plastic type. The lowest AIC value was observed for (Model 4) in the chlorophyll-α and NPP, which included treatment-specific smoothers for both sampling date and plastic concentration, as well as parametric treatment-specific intercepts. Model 2 yielded a slightly lower AIC for ER (ΔAIC = 4.87). However, we selected Model 4 for ER as well to maintain consistency in model structure across metrics and to allow direct ecological comparison of plastic-type-specific effects.

The effects of treatment (plastic type and plastic concentration) on zooplankton biomass and abundance were also analyzed using a generalized additive model (GAM) implemented in the *mgcv* package in R ^64,65^. Plastic concentration was log-transformed prior to analysis, as models using log-transformed concentration had lower AIC values. Zooplankton biomass (µg/L) was estimated by multiplying mean individual dry weights by zooplankton counts (individuals/L). Mean dry weights were calculated using length-to-dry weight regression equations ^66,67^, based on body lengths measured from a subset of individuals collected on days 0, 36, and 97 using ImageJ software ^68^. Taxon-specific regression equations were applied to convert mean lengths to individual dry mass (µg), and these values were then multiplied by the corresponding abundance to estimate population-level biomass.

For both biomass and abundance, a gamma family with a log link function was used to address skewness and ensure normality of residuals before fitting the GAMs. Separate GAMs were applied at each of four time points (0 days, 36 days, 56 days, and 97 days) to avoid overfitting and capture temporal changes. We compared three candidate models using Akaike Information Criterion (AIC), (Model 1) the ‘simplest’ model including only a single global smoother for plastic concentration; (Model 2) a model with plastic type as a parametric term and a shared smoother for plastic concentration and (Model 3) a model including plastic type as a parametric term and plastic-type-specific smoothers for concentration.

For both zooplankton biomass and abundance, Model 3 consistently had the lowest AIC at days 36, 56, and 97, and was therefore selected for these time points. At day 0, Model 1 was the best-fit model for biomass (lowest AIC). For abundance, Model 3 had slightly lower AIC, but as no plastics had been added to ponds, we instead selected Model 1 to avoid overfitting random baseline variation and to maintain parsimony. In this case, the smoother was not significant (p = 0.254), and more complex models offered limited biological interpretability at this pre-treatment time point.

This approach was further supported by PERMANOVA and ANOVA analyses, which revealed no significant differences among treatments in zooplankton community composition (p = 0.928), total abundance (p = 0.43), or Shannon diversity (p = 0.647) at day 0, prior to treatment application.

Model diagnostics were performed using the *gam*.*check* function and ANOVA tables were generated with *anova*.*gam* from the *mgcv* package. Differences between plastic treatments were illustrated by plotting the “difference smooth” using the *plot_difference* function in the *tidymv* package ^69^ (Supplementary Fig. 1, 3, 5-6, 9).

To investigate the main effects of plastic type, plastic concentration, sampling date, and their interactions on eukaryotic and microbial community composition, we conducted a permutational multivariate analysis of variance (PERMANOVA) with the *adonis2* function implemented in the *vegan* package in R ^70^. Chlorophyll-α concentration was included as a covariate in the model to account for variation related to primary productivity. Weighted UniFrac distances were used to account for phylogenetic relationships between communities. To explore temporal variation, separate PERMANOVAs were performed for each sampling date (Day 0, 16, 36, 56, and 94). For these analyses, the Weighted UniFrac distance matrix was subset to include only samples collected on the respective dates. These analyses allowed for a detailed assessment of how plastic type and concentration influenced community composition at specific time points. The models included 1,000 permutations to test for temporal variation in the effects of plastic type and concentration on community structure. Additionally, principal coordinates analysis (PCoA) was performed with the *pcoa* function from the *ape* package ^71^ to visually represent differences in community composition across all sampling dates. A constrained ordination was made from this model using the 2 most predictive axes and their associated species scores. The influence of chlorophyll-⍰ concentrations were represented as a directional vector on the PCoA plots.

To further explore the taxa driving observed patterns in community structure, we conducted biplot analyses by overlaying taxonomic vectors onto previously generated PCoA for days 36, 56, and 94. We applied the *envfit* function from the *vegan* package ^70^ to fit family-level taxa to ordination space based on their relative abundances. Vectors represented the direction of maximum correlation and the strength of association between each taxon and the ordination axes. Only families with significant *envfit* p-values (p < 0.05) were retained, and the ten most abundant among these were displayed for each time point.

To identify the taxa associated with algal blooms in Elastollan tanks, we conducted a differential abundance analysis using 18S rRNA amplicon sequencing data from the last two sampling dates (days 56 and 94). We focused on the six highest plastic concentrations, as they exhibited clear algal blooms. To reduce the influence of low-abundance taxa, we filtered out ASVs with fewer than 10 reads across all samples. Differential abundance testing was performed using the *DESeq2* package ^72^, comparing taxonomic composition between bioplastics and Elastollan tanks. Separate DESeq2 models were run for each sampling date, and taxa with an adjusted p-value (padj) < 0.05 were considered significantly differentially abundant.

## Data availability

The 16S and 18S rRNA amplicon sequencing data are deposited in the NCBI Sequence Read Archive (SRA) under BioProject accession number PRJNA1241327. Processed sequencing data and outputs from initial taxonomic assignments are available at https://github.com/jrdickey9/Bioplastics-1.0. All other datasets generated during the study are available at https://github.com/smorton9292/MP1_Microplastics, and are permanently archived at Zenodo: https://doi.org/10.5281/zenodo.17102858.

## Code availability

All code used for data analysis, statistical modeling, and figure generation is available at https://github.com/smorton9292/MP1_Microplastics. A permanently archived version of the code is also available via Zenodo: https://doi.org/10.5281/zenodo.17102858.

## Acknowledgements

We would like to extend our sincere gratitude to Grant Gustin, Sophia Albee, Lily Raymond, Melissa Santa Cruz, Porsche Noble, and Rachel Tseng for their invaluable assistance with sorting and collecting zooplankton samples, as well as their contributions during fieldwork. We would also like to thank Julia Keum for their assistance in sampling microbial communities. Additionally, we would like to thank Joshua Dominguez and Chris Wall for their insightful discussions and support. All of these contributions were critical to the success of this study.

We also acknowledge the support provided by UCSD Pathways in Biological Sciences Training Program institutional grant from the National Institute of General Medical Sciences (T32 GM133351) which facilitated essential aspects of this research. M.D.B and B.R. were supported by DOE DE-EE0009295. We would like to thank Algenesis Materials for providing TPU FC2.1.

## Author contributions

J.B.S., M.D.B. and S.G.M. conceived the study. B.S.R. and M.D.B. developed and synthesized TPU 181. S.G.M. managed the experiment and coordinated field sampling with contributions from G.V.F., J.R.D., and D.A.L. C.J.S., D.A.L. and J.R.D. processed and curated the sequencing data under the supervision of S.L.J. S.G.M. led the data analysis and figure preparation, with contributions from J.B.S., J.R.D. and G.V.F. S.G.M. wrote the manuscript. All authors contributed to manuscript review and approved the final version.

