## Supplementary information for "Microplastic pollution induces algae blooms in experimental ponds but bioplastics are less harmful"

**Table of Contents**


| *Contents* | *Page #* |
|---|---|
| ***Supplementary Methods*** | |
| Synthesis method for TPU 181 | 2 |
| ***Supplementary Figures*** | |
| Supplementary Figure 1 | 3 |
| Supplementary Figure 2 | 4 |
| Supplementary Figure 3 | 5 |
| Supplementary Figure 4 | 6 |
| Supplementary Figure 5 | 7 |
| Supplementary Figure 6 | 8 |
| Supplementary Figure 7 | 9 |
| Supplementary Figure 8 | 10 |
| Supplementary Figure 9 | 11 |
| Supplementary Figure 10 | 12 |
| Supplementary Figure 11 | 13 |
| Supplementary Figure 12 | 14 |
| Supplementary Figure 13 | 15 |
| Supplementary Figure 14 | 16 |
| Supplementary Figure 15 | 17 |
| Supplementary Figure 16 | 18 |
| ***Supplementary Tables*** | |
| Supplementary Table 1 | 19 |
| Supplementary Table 2 | 20 |
| Supplementary Table 3 | 21 |
| Supplementary Table 4 | 22 |
| Supplementary Table 5 | 23-24 |
| Supplementary Table 6 | 25 |
| Supplementary Table 7 | 26-27 |
| Supplementary Table 8 | 28 |
| Supplementary Table 9 | 29 |

### Supplementary methods

#### Synthesis method for TPU 181

*Algae-Based Polyester-Polyol Synthesis*

The polyester-polyol used for TPU 181 was synthesized following Rajput *et al.* with modifications (Rajput et al. 2022). Briefly, a three-neck round-bottom flask equipped with a Dean–Stark apparatus, reflux condenser, and oil bath was purged and maintained under nitrogen. Commercial azelaic acid (356.4 g, Croda Inc.) and algae-sourced azelaic acid (40.3 g) were combined with 1,3-propanediol (179.2 g) as described in Phung et al. 2020. The mixture was heated to 150–160 °C and gradually raised to 180 °C over one hour. Within the first 4–5 h, water by-product was continuously removed. When approximately 80% of the water was collected, dibutyltin dilaurate (DBTDL, 0.04 wt %) was added. The reaction proceeded for an additional 2–3 days (until desired acid and hydroxyl numbers were reached), with periodic sampling for titration.

*TPU 181 Preparation*

The algae-based polyester-polyol (230 g) and 1,3-propanediol (8.74 g) were dried under vacuum at room temperature for 24 h before polymerization. These were then combined with DBTDL (0.03 wt %) and preheated (75 °C) hexamethylene diisocyanate (6HDI, 42.55 g) in a speed mixer (FlackTek) at 2000 rpm for approximately 1 min. The reaction mixture was poured into molds and cured at 75 °C for 2 days to form TPU sheets.

*Mechanical Property Testing*

Shore A hardness testing was conducted at room temperature. The resulting TPU 181 displayed a Shore A hardness of approximately 97, consistent with typical thermoplastic polyurethanes.

### Supplementary figures

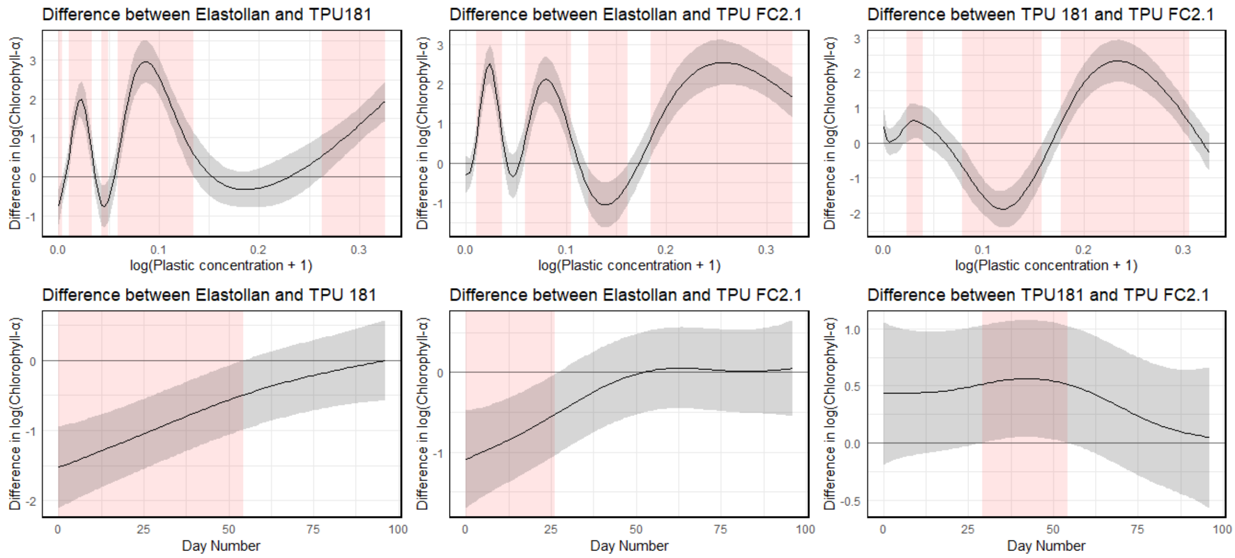


**Supplementary Fig. 1 Significant differences in smooth terms for chlorophyll-α:** Plot showing pairwise differences in the smooth terms for chlorophyll-α between treatments, based on the best fit generalized additive model (GAM). The top row shows differences across plastic concentration, and the bottom row shows differences across day number. Shaded regions represent 95% confidence intervals. Red shaded areas indicate significant divergence between groups.

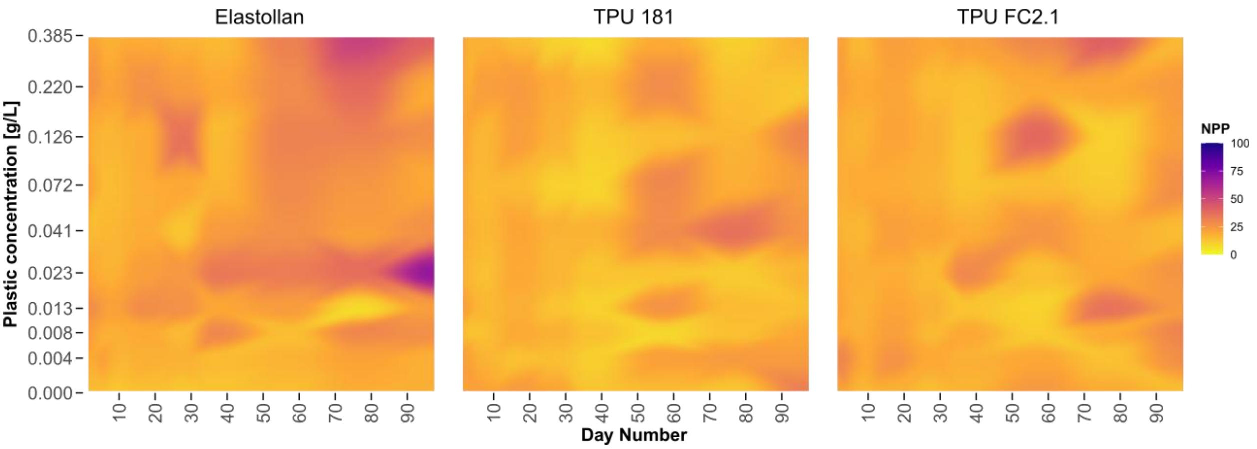


**Supplementary Fig. 2 Net primary production:** Heatmap of net primary production (NPP) for each plastic type across all plastic concentrations over time. Data were interpolated between the nearest NPP measurements to represent a continuous gradient over time.

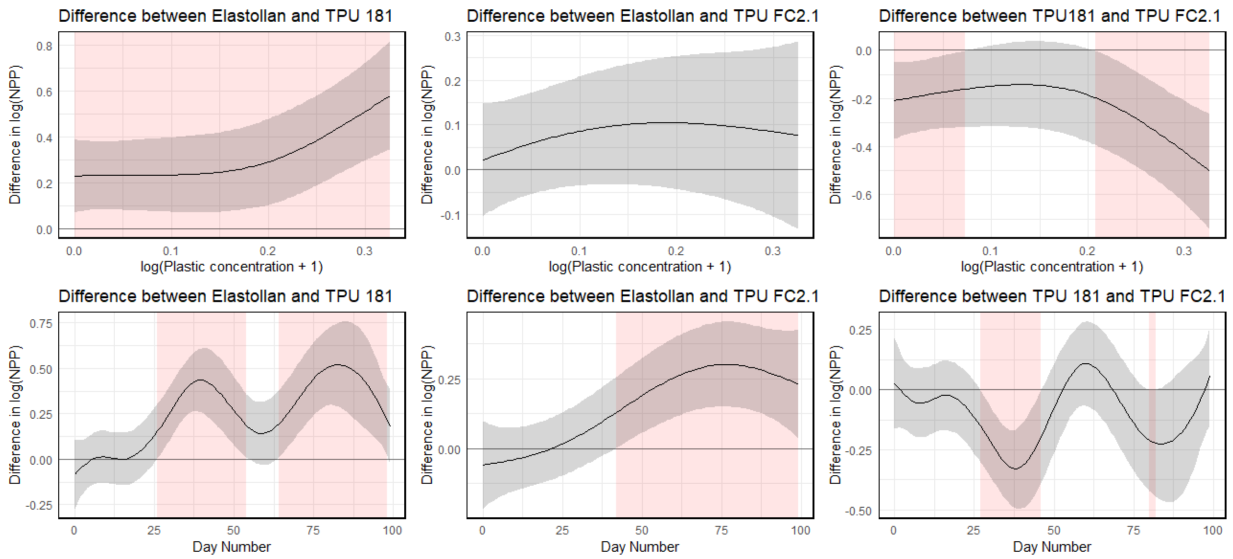


**Supplementary Fig. 3 Significant differences in smooth terms for net primary production:** Plot showing pairwise differences in the smooth terms for net primary production (NPP) between treatments, based on the best fit generalized additive model (GAM). The top row shows differences across plastic concentration, and the bottom row shows differences across day number. Shaded regions represent 95% confidence intervals. Red shaded areas indicate significant divergence between groups.

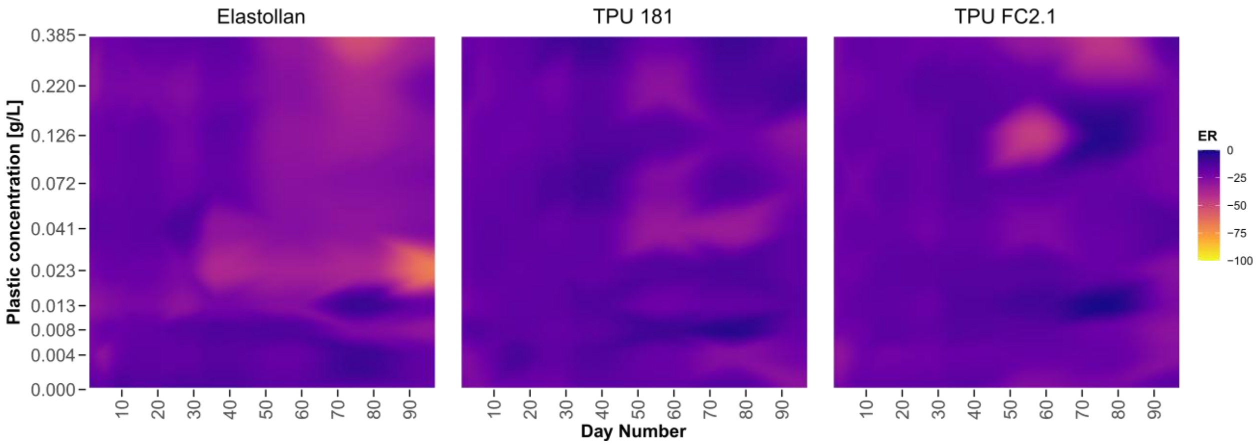


**Supplementary Fig. 4 Ecosystem respiration:** Heatmap of ecosystem respiration (ER) for each plastic type across all plastic concentrations over time. Data were interpolated between the nearest ER measurements to represent a continuous gradient over time.

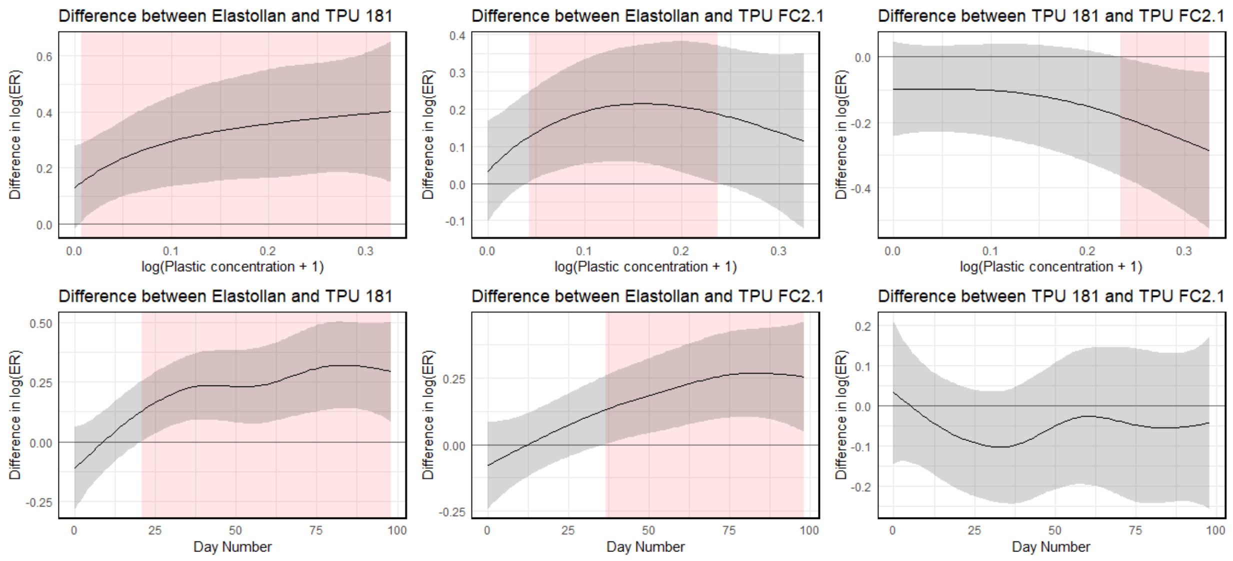


**Supplementary Fig. 5 Significant differences in smooth terms for ecosystem respiration:** Plot showing pairwise differences in the smooth terms for ecosystem respiration (ER) between treatments, based on the best fit generalized additive model (GAM). The top row shows differences across plastic concentration, and the bottom row shows differences across day number. Shaded regions represent 95% confidence intervals. Red shaded areas indicate significant divergence between groups.

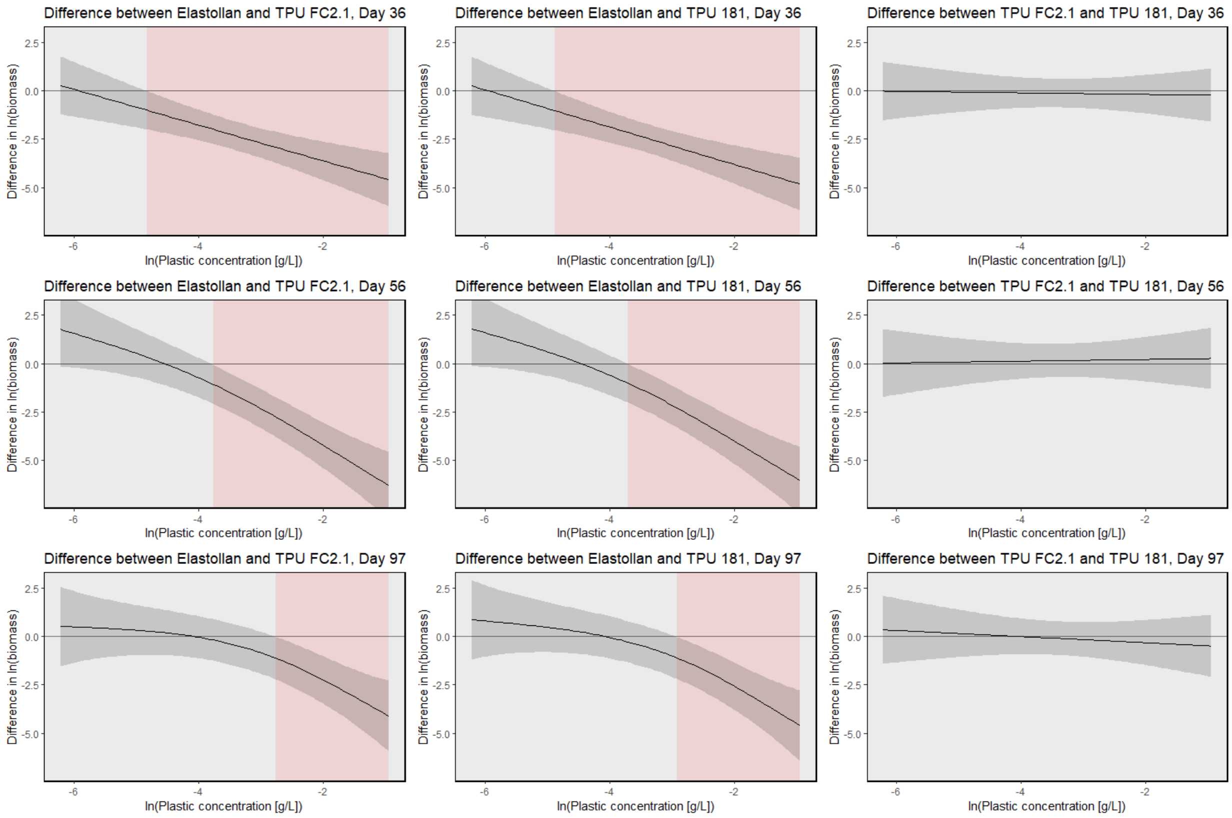


**Supplementary Fig. 6 Significant differences in smooth terms for zooplankton biomass:** Plot showing pairwise differences in the smooth terms for zooplankton biomass between treatments, based on the best fit generalized additive model (GAM). Each row represents a different sampling date across plastic concentration. Shaded regions represent 95% confidence intervals, and red shaded areas indicate significant divergence between groups.

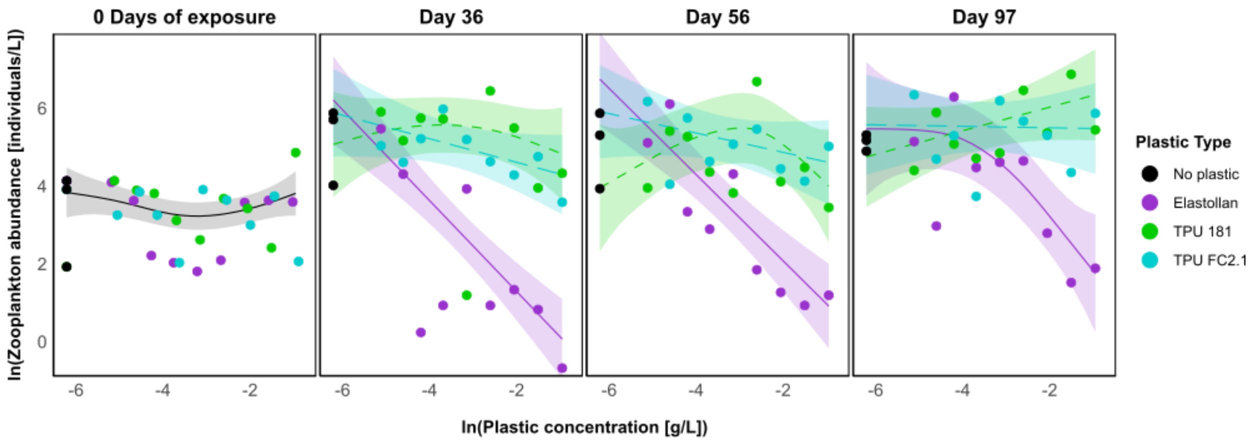


**Supplementary Fig. 7 Zooplankton abundance across plastic concentration and exposure times** Zooplankton abundance [individuals/liter] by plastic concentration [g/L] for each plastic type before application of treatments (left panel, 0 days of exposure) and for 3 dates after the addition of plastics (3 right panels, day 36, 56, and 97). Lines in each plot represent the estimated value across concentrations with a 95% confidence interval.

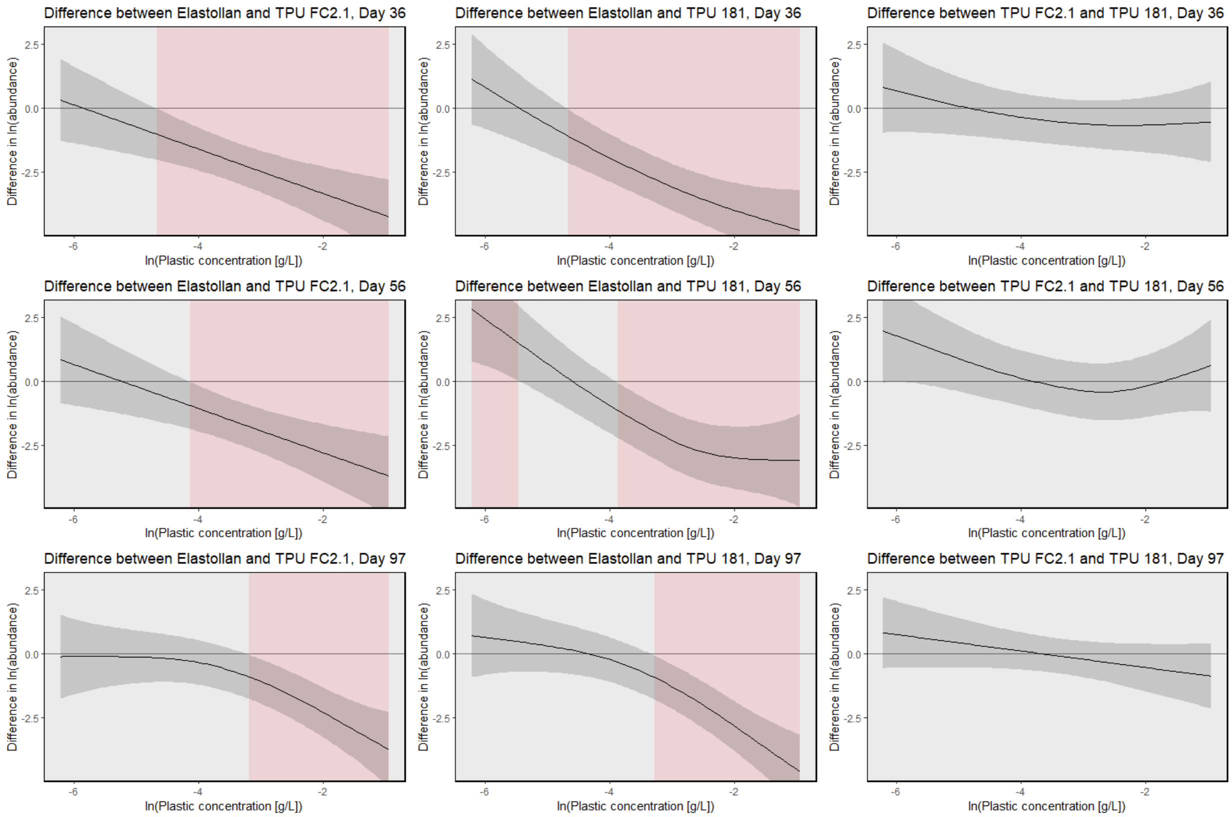


**Supplementary Fig. 8 Significant differences in smooth terms for zooplankton abundance:** Plot showing pairwise differences in the smooth terms for zooplankton abundance between treatments, based on the best fit generalized additive model (GAM). Each row represents a different sampling date across plastic concentration. Shaded regions represent 95% confidence intervals, and red shaded areas indicate significant divergence between groups.

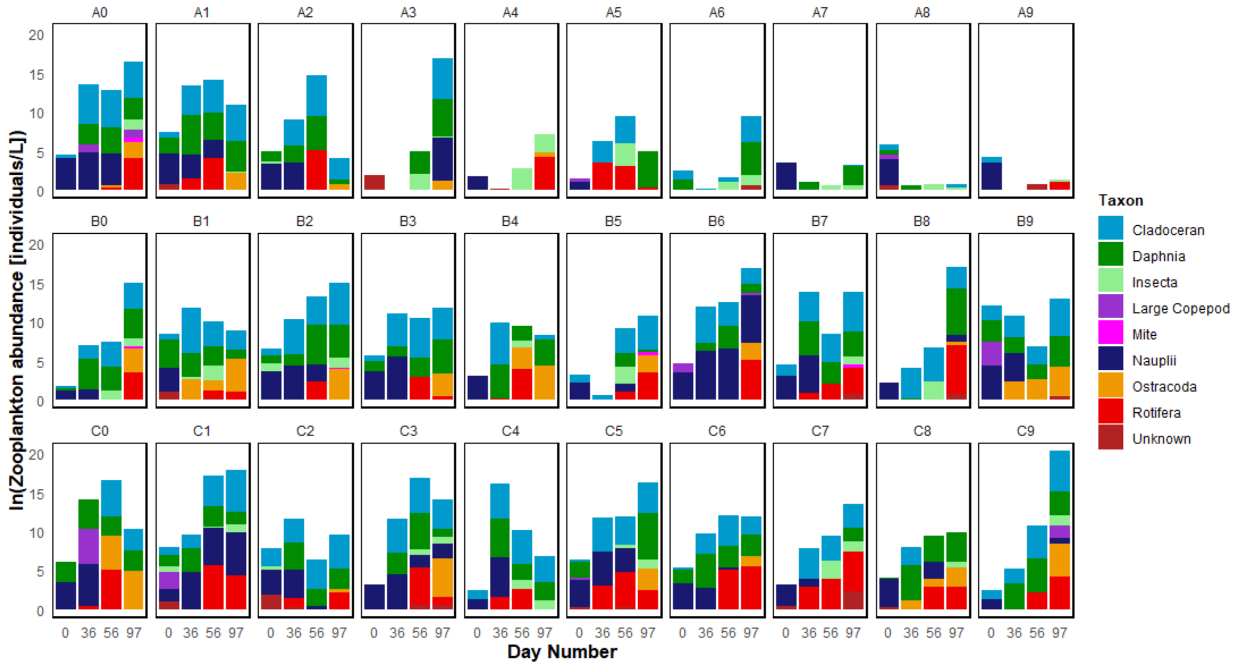


**Supplementary Fig. 9 Zooplankton abundance:** Natural log-transformed zooplankton abundance (ln[individuals/L]) in all experimental ponds on days 0, 36, 56, and 97. Treatments are labeled as A (Elastollan), B (TPU 181), and C (TPU FC2.1), with numbers 0–9 representing increasing experimental concentrations of microplastics. The specific concentrations (g/L) corresponding to labels 0–9 are as follows: 0 = 0.000, 1 = 0.004, 2 = 0.010, 3 = 0.013, 4 = 0.023, 5 = 0.041, 6 = 0.072, 7 = 0.126, 8 = 0.220, and 9 = 0.385.

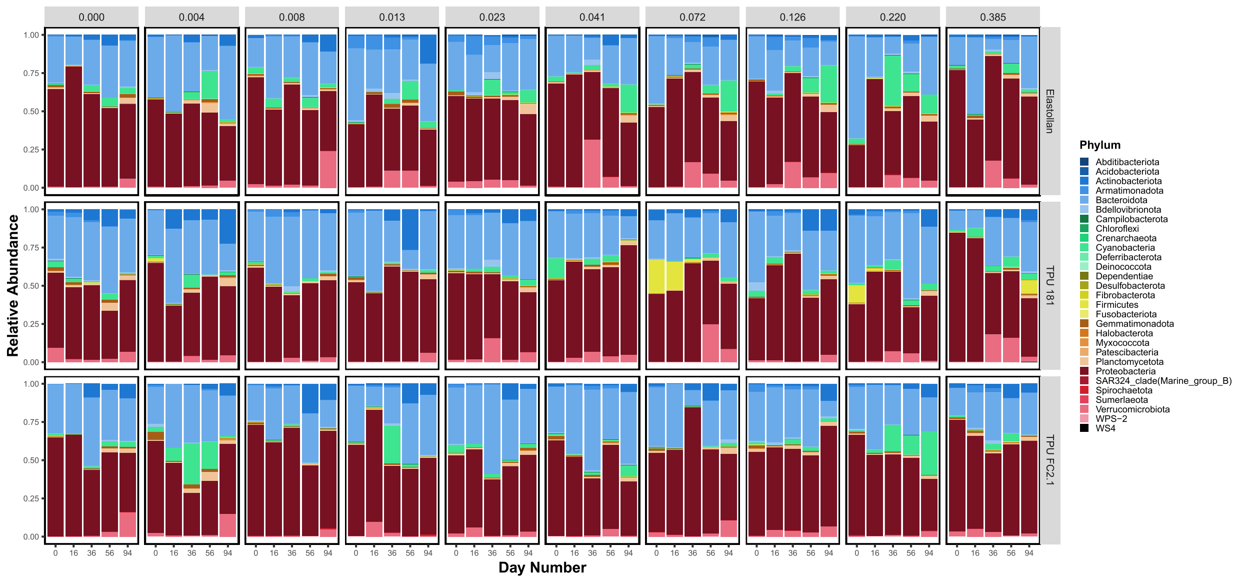


**Supplementary Fig. 10 Relative Abundance of 16S Phyla:** Relative abundance of 16S phyla in all experimental ponds before the application of plastics (0 days of exposure) and for 4 sampling periods after the addition of plastics (day 16, 36, 56, and 94).

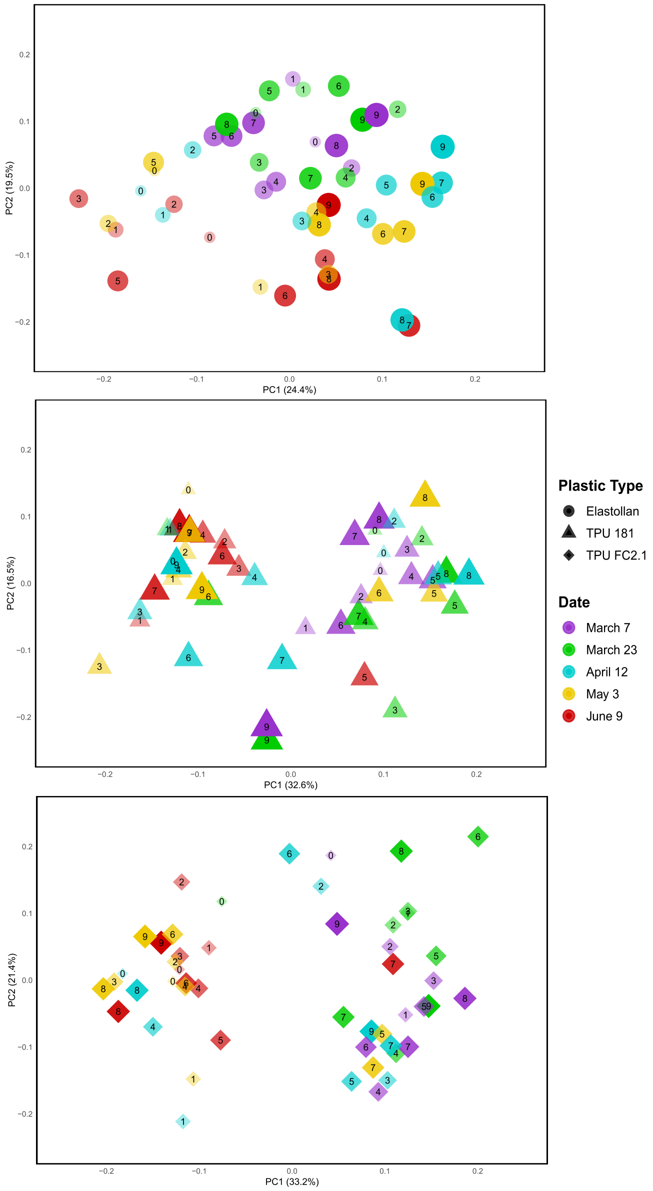


**Supplementary Fig. 11 PCoA 16S all dates:** Principal Coordinates Analysis (PCoA) plots of 16S rRNA data showing the community composition associated with each plastic type across all sampling time points. Colors indicate different sampling time points, and shapes represent plastic type. Numbers within each symbol denote the 10 unique plastic concentrations, including control (0) tanks.

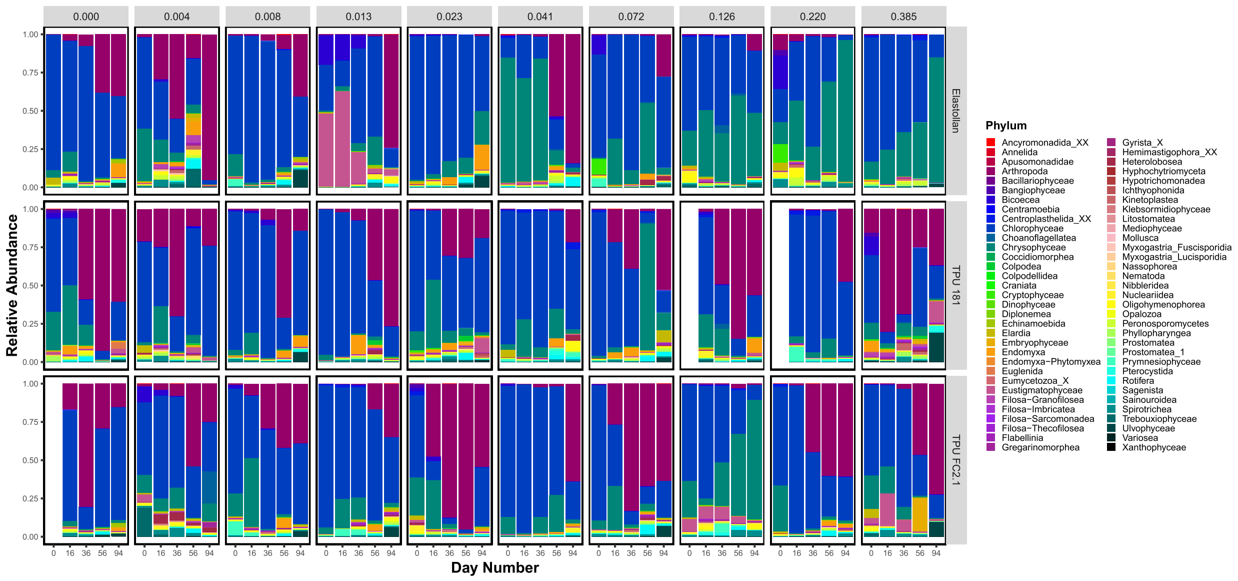


**Supplementary Fig. 12 Relative Abundance of 18S Phyla:** Relative abundance of eukaryotic phyla in all experimental ponds before the application of plastics (0 days of exposure) and for 4 sampling periods after the addition of plastics (day 16, 36, 56, and 94).

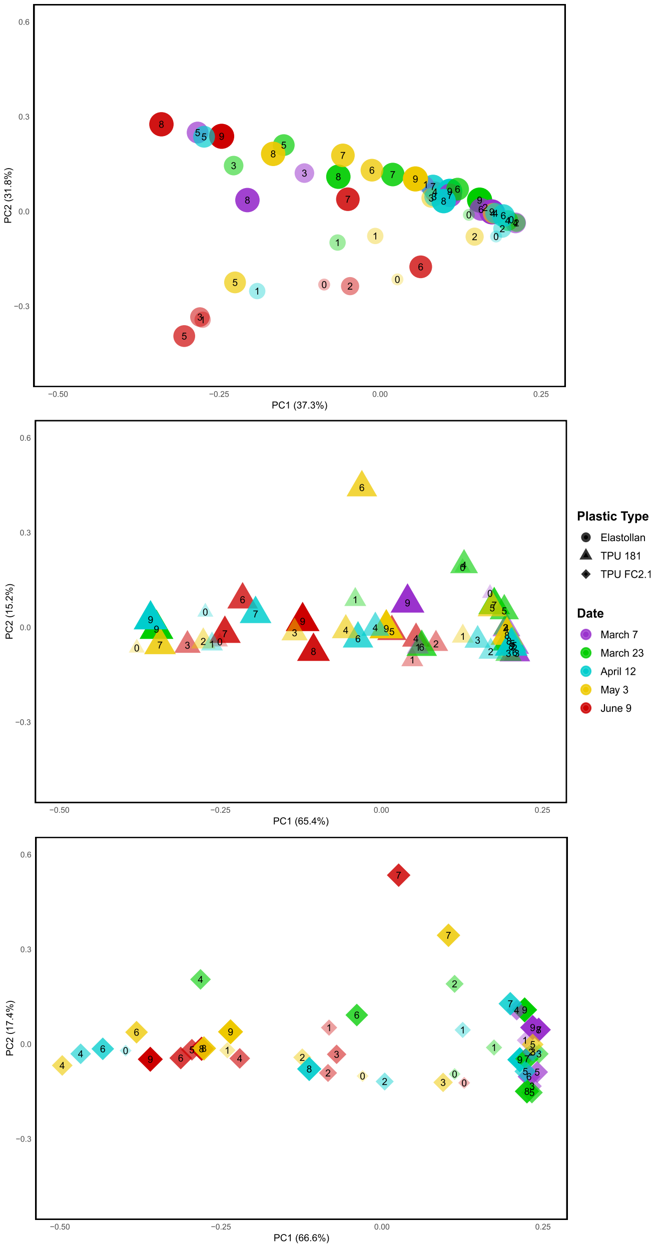


**Supplementary Fig. 13 PCoA 18S all dates:** Principal Coordinates Analysis (PCoA) plots of 18S rRNA data showing the community composition associated with each plastic type across all sampling time points. Colors indicate different sampling time points, and shapes represent plastic type. Numbers within each symbol denote the 10 unique plastic concentrations, including control tanks.

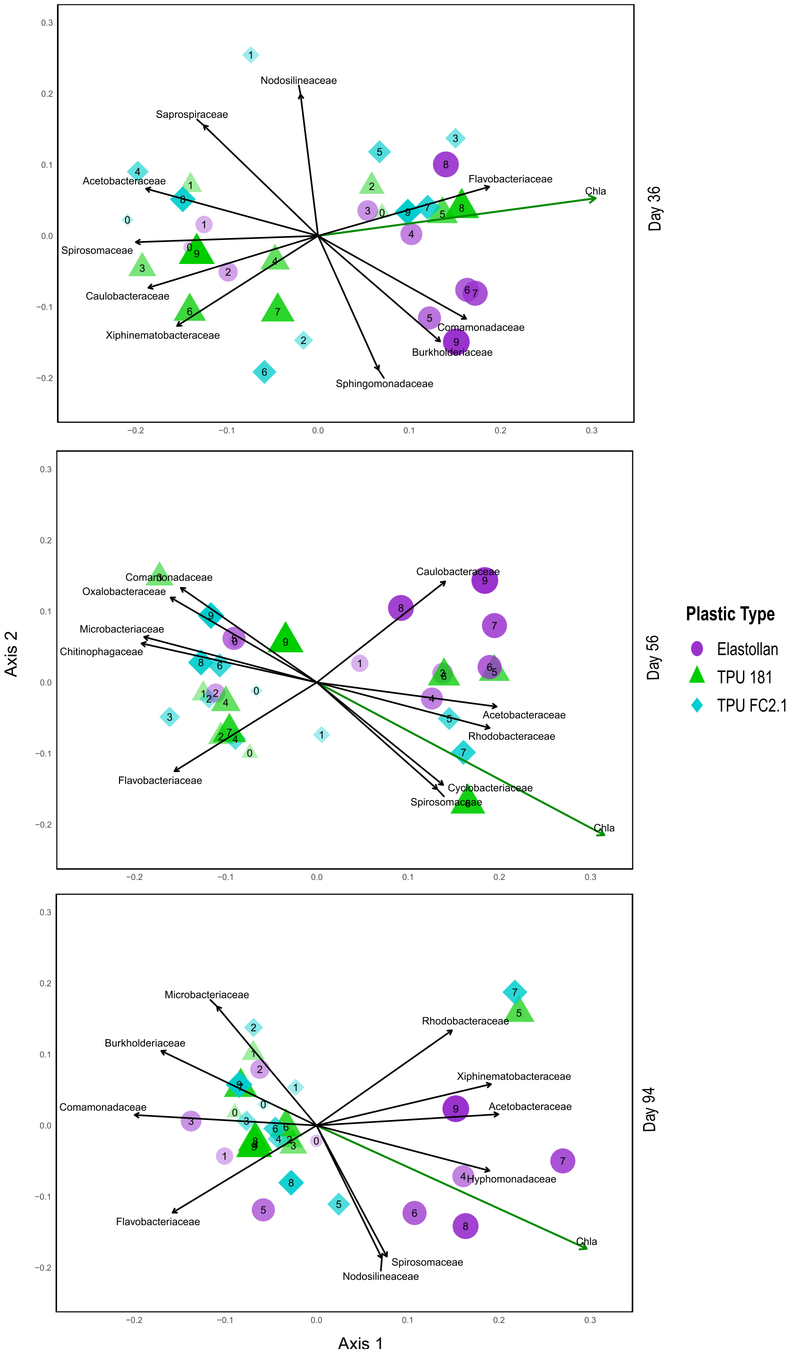


**Supplementary Fig. 14 PCoA ordination with biplots of key 16S taxa**: Principal coordinates analysis (PCoA) ordination plots with biplots overlaid, showing the ten most abundant bacterial families among those significantly correlated with community composition shifts on days 36, 56, and 94 in the 16S dataset. Taxa are represented as vectors indicating their directional influence on community structure across treatments.

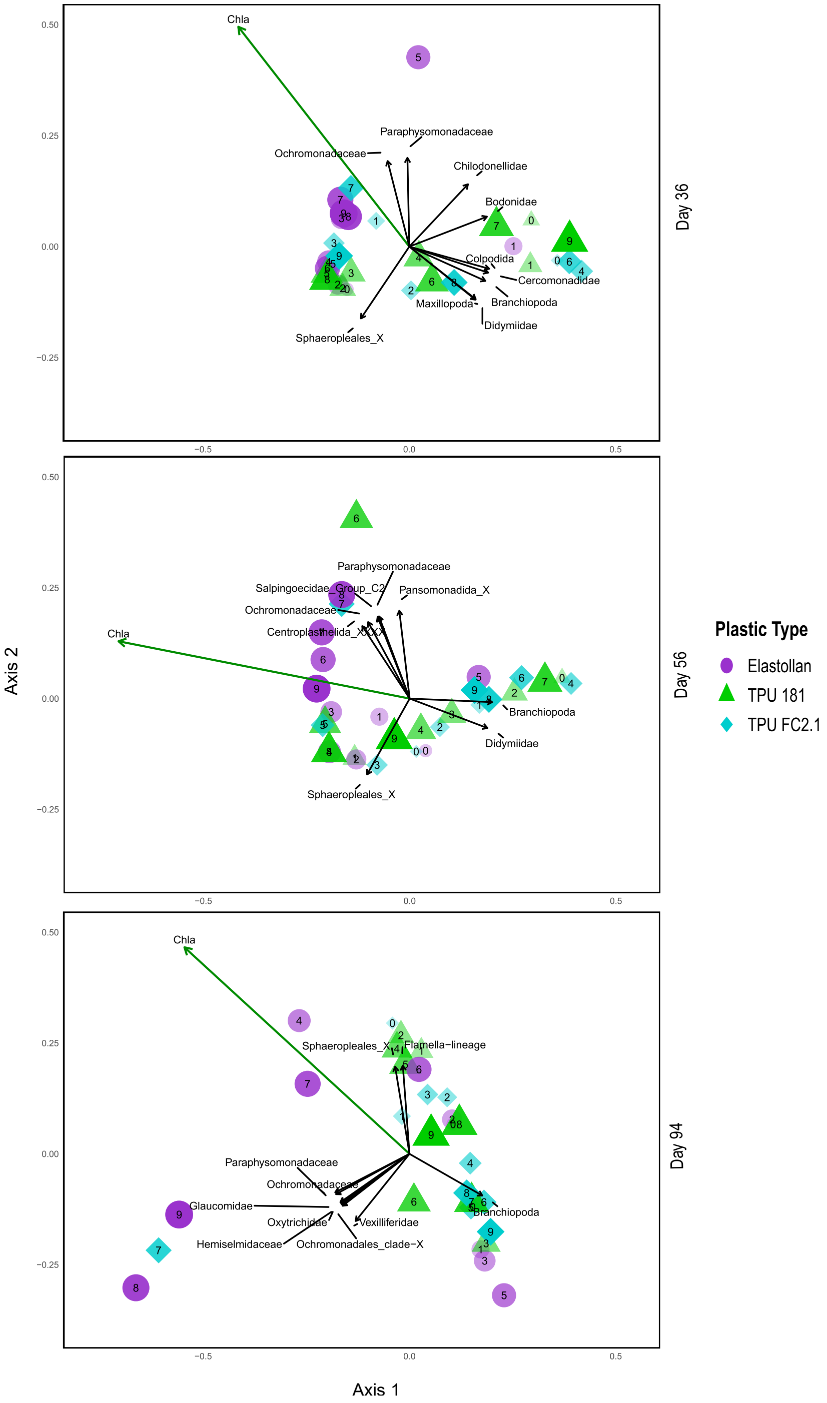


**Supplementary Fig. 15 PCoA ordination with biplots of key 18S taxa**: Principal coordinates analysis (PCoA) ordination plots with biplots overlaid, showing the ten most abundant bacterial families among those significantly correlated with community composition shifts on days 36, 56, and 94 in the 18S dataset. Taxa are represented as vectors indicating their directional influence on community structure across treatments.

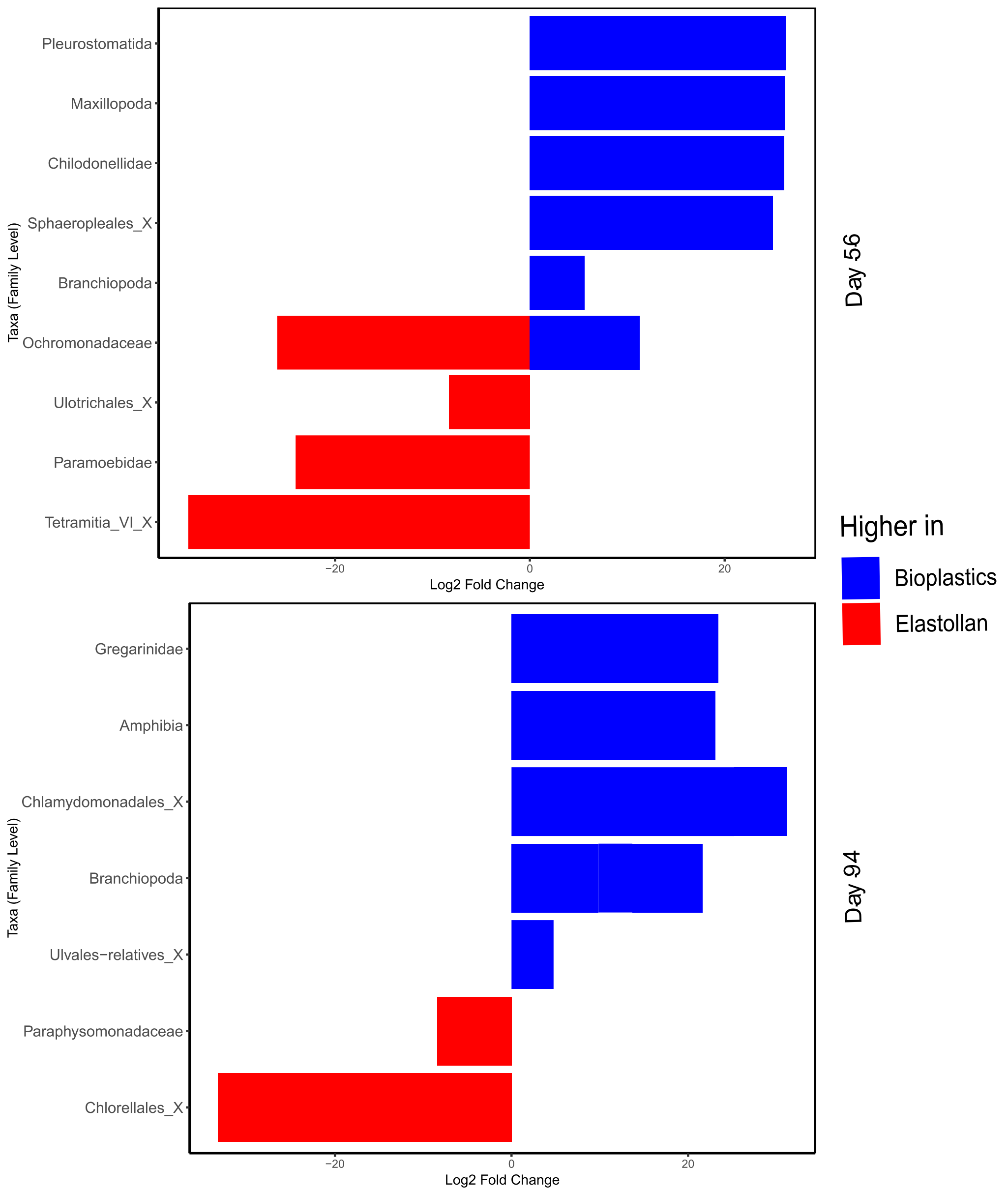


**Supplementary Fig. 16 Differential abundance of enriched 18S taxa on day 56 and 94:** Barplots showing significantly enriched taxa in the 18S datasets, comparing bioplastic and Elastollan tanks from the six highest plastic concentrations on day 56 (top panel) and day 94 (bottom panel). Bars represent taxa with significant $\log_2$ fold changes (padj < 0.05) normalized to control tanks, highlighting differences in community composition between treatment groups.

**Supplementary Table 1 Model selection results for chlorophyll-α, NPP, and ER**

| *Metric* | *Model* | *df* | *AIC* | *ΔAIC* | Adjusted $R^2$ | Deviance Explained (%) |
|---|---|---|---|---|---|---|
| **Chlα** | ~s(day) + s(log_con) | 589.647 | 1750.777 | | | |
| | ~s(day) + s(log_con) + plastic_type | 587.548 | 1716.672 | | | |
| | ~s(day, by=plastic_type) + s(log_con, by=plastic_type) | 571.414 | 1505.872 | | | |
| | ~s(day, by=plastic_type) + s(log_con, by=plastic_type) + plastic_type | 568.524 | 1446.793 | -303.984 | 0.547 | 56.9 |
| **NPP** | ~s(day) + s(log_con) | 10.333 | 686.415 | | | |
| | ~s(day) + s(log_con) + plastic_type | 12.505 | 660.691 | | | |
| | ~s(day, by=plastic_type) + s(log_con, by=plastic_type) | 20.210 | 675.805 | | | |
| | ~s(day, by=plastic_type) + s(log_con, by=plastic_type) + plastic_type | 22.572 | 648.564 | -37.850 | 0.147 | 17.0 |
| **ER** | ~s(day) + s(log_con) | 10.028 | 361.444 | | | |
| | ~s(day) + s(log_con) + plastic_type | 12.210 | 343.972 | | | |
| | ~s(day, by=plastic_type) + s(log_con, by=plastic_type) | 16.135 | 366.300 | | | |
| | ~s(day, by=plastic_type) + s(log_con, by=plastic_type) + plastic_type | 18.492 | 348.839 | -12.605 | 0.113 | 14.1 |

*Metric* refers to the ecological response variable analyzed (chlorophyll-α, NPP, or ER). *Model* indicates the fixed effects structure of each candidate model. *df* is the number of estimated degrees of freedom. *AIC* is Akaike's Information Criterion value, and *ΔAIC* is the difference in AIC between the simplest model and the selected model. *Adjusted* $R^2$ represents the proportion of variance explained by the model, adjusted for model complexity. *Deviance Explained (%)* indicates the percentage of deviance in the response variable accounted for by the model.

**Supplementary Table 2 Generalized additive model (GAM) results for chlorophyll-α, NPP, and ER across plastic type**

| *Metric* | Model | *Estimate* | *Std. Error* | *t-value* | *df/edf* | *Ref.df* | *F* | *p-value* |
|---|---|---|---|---|---|---|---|---|
| **Chlα** | ~s(day, by=plastic_type) + s(log_con, by=plastic_type) | | | | | | | |
| | Parametric: Plastic Type Elastollan (*intercept*) | 3.187 | 0.056 | 57.310 | | | | <0.0001 |
| | Parametric: Plastic Type TPU 181 | -0.567 | 0.079 | -7.213 | | | | <0.0001 |
| | Parametric: Plastic Type TPU FC2.1 | -0.509 | 0.079 | -6.482 | | | | <0.0001 |
| | s(day) : Elastollan | | | | 1.317 | 1.567 | 5.982 | 0.0047 |
| | s(day): TPU 181 | | | | 1.750 | 2.181 | 13.056 | <0.0001 |
| | s(day) : TPU FC2.1 | | | | 2.922 | 3.629 | 4.980 | 0.0019 |
| | s(log_con) : Elastollan | | | | 7.171 | 7.719 | 28.958 | <0.0001 |
| | s(log_con): TPU 181 | | | | 7.168 | 7.717 | 25.806 | <0.0001 |
| | s(log_con) : TPU FC2.1 | | | | 7.147 | 7.697 | 24.344 | <0.0001 |
| **NPP** | ~s(day, by=plastic_type) + s(log_con, by=plastic_type) + plastic_type | | | | | | | |
| | Parametric: Plastic Type Elastollan (*intercept*) | 3.058 | 0.025 | 120.283 | | | | <0.0001 |
| | Parametric: Plastic Type TPU 181 | -0.200 | 0.036 | -5.570 | | | | <0.0001 |
| | Parametric: Plastic Type TPU FC2.1 | -0.102 | 0.036 | -2.839 | | | | 0.0046 |
| | s(day) : Elastollan | | | | 2.602 | 3.184 | 8.447 | <0.0001 |
| | s(day): TPU 181 | | | | 5.907 | 6.729 | 6.699 | <0.0001 |
| | s(day) : TPU FC2.1 | | | | 2.428 | 2.974 | 1.937 | 0.1113 |
| | s(log_con) : Elastollan | | | | 1.050 | 1.098 | 5.206 | 0.0182 |
| | s(log_con): TPU 181 | | | | 2.227 | 2.682 | 2.017 | 0.0815 |
| | s(log_con) : TPU FC2.1 | | | | 1.562 | 1.904 | 2.141 | 0.1768 |
| **ER** | ~s(day, by=plastic_type) + s(log_con, by=plastic_type) + plastic_type | | | | | | | |
| | Parametric: Plastic Type Elastollan (*intercept*) | 3.093 | 0.029 | 107.664 | | | | <0.0001 |
| | Parametric: Plastic Type TPU 181 | -0.185 | 0.041 | -4.557 | | | | <0.0001 |
| | Parametric: Plastic Type TPU FC2.1 | -0.115 | 0.041 | -2.842 | | | | 0.0047 |
| | s(day) : Elastollan | | | | 1.770 | 2.178 | 5.692 | 0.00353 |
| | s(day): TPU 181 | | | | 3.172 | 3.857 | 2.960 | 0.01833 |
| | s(day) : TPU FC2.1 | | | | 2.305 | 2.815 | 1.854 | 0.15563 |
| | s(log_con) : Elastollan | | | | 2.211 | 2.663 | 3.458 | 0.02933 |
| | s(log_con): TPU 181 | | | | 1.619 | 1.976 | 1.066 | 0.38306 |
| | s(log_con) : TPU FC2.1 | | | | 1.001 | 1.002 | 0.966 | 0.32674 |

*Metric* refers to the ecological response variable analyzed. *Model* indicates the predictor term (e.g., plastic type or smoothing term). *Estimate* is the estimated effect size for parametric terms. *Std. Error* is the standard error of the estimate. *t-value* is the test statistic for parametric terms. *df/edf* is the degrees of freedom (df) for parametric terms or estimated degrees of freedom (edf) for smooth terms. *Ref.df* is the reference degrees of freedom used to calculate the F-statistic. *F* is the F-statistic for smooth terms. *p-value* indicates the significance of the model term.

**Supplementary Table 3 Model selection results for zooplankton biomass and abundances**

| *Metric* | *Time* | *Model* | *df* | *AIC* | *Δ AIC* | Adjusted $R^2$ | Deviance Explained (%) |
|---|---|---|---|---|---|---|---|
| **Biomass** | 0 Days | ~ s(log.con) | 7.80 | 281.9873 | 0 | 0.659 | 45.4 |
| | | ~ plastic_type + s(log.con) | 9.23 | 283.8462 | | | |
| | | ~ plastic_type + s(log.con, by=plastic_type) | 9.43 | 284.0886 | | | |
| | Day 36 | ~ s(log.con) | 3.00 | 381.2262 | | | |
| | | ~ plastic_type + s(log.con) | 5.00 | 377.2943 | | | |
| | | ~ plastic_type + s(log.con, by=plastic_type) | 7.00 | 374.2129 | -7.0133 | 0.191 | 40.2 |
| | Day 56 | ~ s(log.con) | 3.00 | 373.8640 | | | |
| | | ~ plastic_type + s(log.con) | 5.00 | 376.0841 | | | |
| | | ~ plastic_type + s(log.con, by=plastic_type) | 8.06 | 366.2181 | -7.6459 | -1.940 | 43.2 |
| | Day 97 | ~ s(log.con) | 3.00 | 420.2585 | | | |
| | | ~ plastic_type + s(log.con) | 5.00 | 422.8167 | | | |
| | | ~ plastic_type + s(log.con, by=plastic_type) | 8.55 | 417.2067 | -3.0518 | -0.035 | 34.9 |
| **Abundance** | 0 Days | ~ s(log.con) | 4.83 | 269.7680 | 0 | 0.078 | 14.5 |
| | | ~ plastic_type + s(log.con) | 6.87 | 272.4354 | | | |
| | | ~ plastic_type + s(log.con, by=plastic_type) | 13.29 | 265.1669 | | | |
| | Day 36 | ~ s(log.con) | 3.00 | 357.6869 | | | |
| | | ~ plastic_type + s(log.con) | 5.00 | 348.9697 | | | |
| | | ~ plastic_type + s(log.con, by=plastic_type) | 8.01 | 340.4372 | -17.2497 | 0.257 | 58.3 |
| | Day 56 | ~ s(log.con) | 3.00 | 362.4874 | | | |
| | | ~ plastic_type + s(log.con) | 5.00 | 360.5040 | | | |
| | | ~ plastic_type + s(log.con, by=plastic_type) | 8.93 | 346.8757 | -15.6117 | -0.450 | 60.4 |
| | Day 97 | ~ s(log.con) | 3.00 | 390.1397 | | | |
| | | ~ plastic_type + s(log.con) | 5.00 | 389.9740 | | | |
| | | ~ plastic_type + s(log.con, by=plastic_type) | 8.70 | 377.8518 | -12.2879 | 0.098 | 52.4 |

*Metric* refers to the zooplankton response variable analyzed (biomass or abundance). *Time* indicates the sampling period included in the model. *Model* describes the structure of each candidate model. *df* is the number of estimated degrees of freedom. *AIC* is Akaike's Information Criterion value, and *ΔAIC* is the difference in AIC between the simplest model and the selected model. *Adjusted* $R^2$ represents the proportion of variance explained by the model, adjusted for model complexity. *Deviance Explained (%)* indicates the percentage of deviance in the response variable accounted for by the model.

**Supplementary Table 4 Generalized additive model (GAM) results for zooplankton biomass and abundance across sampling periods**

| *Metric* | *Time* | *Effect* | *df/edf* | *Ref.df* | *F* | *p-value* |
|---|---|---|---|---|---|---|
| **Biomass** | 0 Days | s(log.con) | 4.792 | 5.807 | 2.918 | 0.0262 |
| | Day 36 | plastic_type + s(log.con, by = plastic_type) | 2 | | 26.620 | <0.0001 |
| | | s(log.con) : Elastollan | 1 | 1 | 54.150 | <0.0001 |
| | | s(log.con): TPU 181 | 1 | 1 | 2.670 | 0.1153 |
| | | s(log.con) : TPU FC2.1 | 1 | 1 | 3.510 | 0.0732 |
| | Day 56 | plastic_type + s(log.con, by = plastic_type) | 2 | | 12.710 | 0.0002 |
| | | s(log.con) : Elastollan | 1.654 | 2.057 | 36.595 | <0.0001 |
| | | s(log.con) : TPU 181 | 1 | 1 | 0.579 | 0.4550 |
| | | s(log.con) : TPU FC2.1 | 1 | 1 | 0.282 | 0.6010 |
| | Day 97 | plastic_type + s(log.con, by = plastic_type) | 2 | | 4.166 | 0.0286 |
| | | s(log.con) : Elastollan | 2.042 | 2.545 | 9.428 | 0.0007 |
| | | s(log.con) : TPU 181 | 1 | 1 | 0.915 | 0.3487 |
| | | s(log.con) : TPU FC2.1 | 1 | 1 | 0.024 | 0.8788 |
| **Aubundance** | 0 Days | s(log.con) | 2.273 | 2.825 | 1.777 | 0.2540 |
| | Day 36 | plastic_type + s(log.con, by = plastic_type) | 2 | | 21.320 | <0.0001 |
| | | s(log.con) : Elastollan | 1 | 1 | 42.889 | <0.0001 |
| | | s(log.con): TPU 181 | 1.621 | 2.012 | 0.734 | 0.4960 |
| | | s(log.con) : TPU FC2.1 | 1 | 1 | 2.881 | 0.1030 |
| | Day 56 | plastic_type + s(log.con, by = plastic_type) | 2 | | 7.624 | 0.0029 |
| | | s(log.con) : Elastollan | 1.002 | 1.003 | 34.055 | <0.0001 |
| | | s(log.con) : TPU 181 | 2.358 | 2.927 | 1.569 | 0.2700 |
| | | s(log.con) : TPU FC2.1 | 1 | 1 | 1.717 | 0.2030 |
| | Day 97 | plastic_type + s(log.con, by = plastic_type) | 2 | | 9.068 | 0.0013 |
| | | s(log.con) : Elastollan | 2.173 | 2.704 | 10.358 | 0.0003 |
| | | s(log.con) : TPU 181 | 1 | 1 | 3.801 | 0.0502 |
| | | s(log.con) : TPU FC2.1 | 1 | 1 | 0.015 | 0.9047 |

*Metric* refers to the zooplankton response variable analyzed (biomass or abundance). *Time* indicates the sampling period included in the model. *Effect* denotes the model term (parametric or smooth) being evaluated. *df/edf* is the degrees of freedom (df) for parametric terms or estimated degrees of freedom (edf) for smooth terms. *Ref.df* is the reference degrees of freedom used in calculating the F-statistic. *F* is the F-statistic for the effect, and *p-value* indicates the significance of the model term.

**Supplementary Table 5 Significant pairwise comparisons of zooplankton biomass between plastic treatments across concentrations and sampling dates**

| *Day Number* | *Plastic Concentration (g/L)* | *Comparison* | *Estimate ± SE* | *t-ratio* | *p-value* |
|---|---|---|---|---|---|
| Day 36 | 0.008 | Elastollan - TPU 181 | -1.293 ± 0.473 | -2.736 | 0.0299 |
| | 0.008 | Elastollan - TPU 2.1 | -1.212 ± 0.473 | -2.565 | 0.0433 |
| | 0.013 | Elastollan - TPU 181 | -1.683 ± 0.422 | -3.993 | 0.0015 |
| | 0.013 | Elastollan - TPU 2.1 | -1.586 ± 0.422 | -3.762 | 0.0027 |
| | 0.023 | Elastollan - TPU 181 | -2.176 ± 0.383 | -5.678 | <0.0001 |
| | 0.023 | Elastollan - TPU 2.1 | -2.058 ± 0.383 | -5.371 | <0.0001 |
| | 0.041 | Elastollan - TPU 181 | -2.698 ± 0.382 | -7.056 | <0.0001 |
| | 0.041 | Elastollan - TPU 2.1 | -2.558 ± 0.382 | -6.690 | <0.0001 |
| | 0.072 | Elastollan - TPU 181 | -3.221 ± 0.423 | -7.613 | <0.0001 |
| | 0.072 | Elastollan - TPU 2.1 | -3.059 ± 0.423 | -7.231 | <0.0001 |
| | 0.126 | Elastollan - TPU 181 | -3.749 ± 0.496 | -7.560 | <0.0001 |
| | 0.126 | Elastollan - TPU 2.1 | -3.565 ± 0.496 | -7.189 | <0.0001 |
| | 0.220 | Elastollan - TPU 181 | -4.279 ± 0.589 | -7.260 | <0.0001 |
| | 0.220 | Elastollan - TPU 2.1 | -4.073 ± 0.589 | -6.911 | <0.0001 |
| | 0.385 | Elastollan - TPU 181 | -4.814 ± 0.696 | -6.916 | <0.0001 |
| | 0.385 | Elastollan - TPU 2.1 | -4.586 ± 0.696 | -6.588 | <0.0001 |
| Day 56 | 0.041 | Elastollan - TPU 181 | -1.895 ± 0.509 | -3.723 | 0.0030 |
| | 0.041 | Elastollan - TPU 2.1 | -2.059 '± 0.509 | -4.044 | 0.0014 |
| | 0.072 | Elastollan - TPU 181 | -2.853 ± 0.534 | -5.344 | 0.0001 |
| | 0.072 | Elastollan - TPU 2.1 | -3.041 ± 0.534 | -5.696 | <0.0001 |
| | 0.126 | Elastollan - TPU 181 | -3.89 ± 0.593 | -6.563 | <0.0001 |
| | 0.126 | Elastollan - TPU 2.1 | -4.103 ± 0.593 | -6.922 | <0.0001 |
| | 0.220 | Elastollan - TPU 181 | -4.956 ± 0.703 | -7.050 | <0.0001 |
| | 0.220 | Elastollan - TPU 2.1 | -5.193 ± 0.703 | -7.387 | <0.0001 |
| | 0.385 | Elastollan - TPU 181 | -6.036 ± 0.884 | -6.824 | <0.0001 |
| | 0.385 | Elastollan - TPU 2.1 | -6.298 ± 0.884 | -7.120 | <0.0001 |
| Day 97 | 0.072 | Elastollan - TPU 181 | -1.547 ± 0.568 | -2.723 | 0.0314 |
| | 0.126 | Elastollan - TPU 181 | -2.467 ± 0.616 | -4.007 | 0.0015 |
| | 0.126 | Elastollan - TPU 2.1 | -2.15 ± 0.616 | -3.492 | 0.0054 |
| | 0.220 | Elastollan - TPU 181 | -3.512 ± 0.719 | -4.882 | 0.0002 |
| | 0.220 | Elastollan - TPU 2.1 | -3.107 ± 0.719 | -4.320 | 0.0007 |
| | 0.385 | Elastollan - TPU 181 | -4.605 ± 0.931 | -4.949 | 0.0002 |
| | 0.385 | Elastollan - TPU 2.1 | -4.113 ± 0.931 | -4.420 | 0.0006 |

*Day Number* indicates the sampling period. *Plastic Concentration (g/L)* refers to the concentration of plastic associated with each comparison. *Comparison* specifies the pair of plastic treatments being compared. *Estimate ± SE* is the difference in estimated zooplankton biomass between the two treatments, reported with the standard error. *t-ratio* is the test statistic from the pairwise comparison. *p-value* indicates the significance of the comparison.

**Supplementary Table 6 16S and 18S global PERMANOVA results**

| *Metric* | *Effect* | *df* | *SumofSqs* | *R²* | *F* | *p-value* |
|---|---|---|---|---|---|---|
| **16S** | plastic_type | 2 | 0.2032 | 0.02998 | 2.8401 | 0.00400 |
| | plastic_concentration | 1 | 0.0780 | 0.01151 | 2.1807 | 0.02997 |
| | date | 4 | 1.0643 | 0.15706 | 7.4384 | 0.00100 |
| | chlα | 1 | 0.3345 | 0.04936 | 9.3510 | 0.00100 |
| | plastic_type:plastic_concentration | 2 | 0.151 | 0.22280 | 2.1108 | 0.01399 |
| | plastic_type:date | 8 | 0.3638 | 0.05368 | 1.2712 | 0.09391 |
| | plastic_concentration:date | 4 | 0.1072 | 0.15820 | 0.7493 | 0.85315 |
| | plastic_type:plastic_concentration_date | 8 | 0.2178 | 0.03214 | 0.7611 | 0.92408 |
| **18S** | plastic_type | 2 | 0.3852 | 0.04204 | 4.1079 | 0.00300 |
| | plastic_concentration | 1 | 0.0897 | 0.00979 | 1.9141 | 0.11688 |
| | date | 4 | 1.5304 | 0.16703 | 8.1611 | 0.00100 |
| | chlα | 1 | 0.4952 | 0.05405 | 10.5625 | 0.00100 |
| | plastic_type:plastic_concentration | 2 | 0.2396 | 0.02615 | 2.5551 | 0.03497 |
| | plastic_type:date | 8 | 0.3617 | 0.03948 | 0.9645 | 0.50350 |
| | plastic_concentration:date | 4 | 0.1797 | 0.01962 | 0.9584 | 0.46154 |
| | plastic_type:plastic_concentration_date | 8 | 0.4427 | 0.04832 | 1.1803 | 0.25475 |

*Metric* refers to the dataset analyzed (16S or 18S). *Effect* indicates the explanatory variable tested. *df* is the degrees of freedom for each effect. *SumofSqs* is the sum of squares associated with the effect. $R^2$ represents the proportion of variance explained by the effect. *F* is the F-statistic from the PERMANOVA test. *p-value* indicates the significance of the effect.

**Supplementary Table 7 Date specific 16S and 18S PERMANOVA results**

| *Metric* | *Time* | *Effect* | *df* | *SumofSqs* | *R²* | *F* | *p-value* |
|---|---|---|---|---|---|---|---|
| **16S** | 0 Days | plastic_type | 2 | 0.0813 | 0.0848 | 1.2438 | 0.1848 |
| | | plastic_concentration | 1 | 0.0374 | 0.0391 | 1.1455 | 0.2967 |
| | | chlα | 1 | 0.4486 | 0.0468 | 1.3729 | 0.2058 |
| | | plastic_type:plastic_concentration | 2 | 0.0434 | 0.0453 | 0.6643 | 0.8561 |
| | Day 16 | plastic_type | 2 | 0.1167 | 0.1037 | 1.6519 | 0.0839 |
| | | plastic_concentration | 1 | 0.0168 | 0.0149 | 0.4757 | 0.8801 |
| | | chlα | 1 | 0.1105 | 0.0982 | 3.1284 | 0.0060 |
| | | plastic_type:plastic_concentration | 2 | 0.0685 | 0.0609 | 0.9705 | 0.4525 |
| | Day 36 | plastic_type | 2 | 0.1523 | 0.1111 | 1.9516 | 0.0370 |
| | | plastic_concentration | 1 | 0.0533 | 0.0389 | 1.3658 | 0.2068 |
| | | chlα | 1 | 0.1846 | 0.1347 | 4.7313 | 0.0010 |
| | | plastic_type:plastic_concentration | 2 | 0.0833 | 0.0608 | 1.0673 | 0.4086 |
| | Day 56 | plastic_type | 2 | 0.1388 | 0.1179 | 2.3266 | 0.0220 |
| | | plastic_concentration | 1 | 0.0538 | 0.0455 | 1.8030 | 0.0969 |
| | | chlα | 1 | 0.2077 | 0.1764 | 6.9608 | 0.0010 |
| | | plastic_type:plastic_concentration | 2 | 0.0910 | 0.0773 | 1.5253 | 0.1319 |
| | Day 94 | plastic_type | 2 | 0.1104 | 0.0964 | 1.4821 | 0.1039 |
| | | plastic_concentration | 1 | 0.0271 | 0.0251 | 0.7717 | 0.6134 |
| | | chlα | 1 | 0.0570 | 0.0527 | 1.6205 | 0.1439 |
| | | plastic_type:plastic_concentration | 2 | 0.0846 | 0.0783 | 1.2036 | 0.2627 |
| **18S** | 0 Days | plastic_type | 2 | 0.0842 | 0.0976 | 1.3657 | 0.2058 |
| | | plastic_concentration | 1 | 0.0383 | 0.0444 | 1.2421 | 0.2797 |
| | | chlα | 1 | 0.0730 | 0.0846 | 2.3678 | 0.0799 |
| | | plastic_type:plastic_concentration | 2 | 0.0510 | 0.0591 | 0.8270 | 0.4565 |
| | Day 16 | plastic_type | 2 | 0.0510 | 0.0379 | 0.5947 | 0.7892 |
| | | plastic_concentration | 1 | 0.0138 | 0.0102 | 0.3208 | 0.8621 |
| | | chlα | 1 | 0.1263 | 0.0939 | 2.9455 | 0.0340 |
| | | plastic_type:plastic_concentration | 2 | 0.1671 | 0.1244 | 1.9495 | 0.1079 |
| | Day 36 | plastic_type | 2 | 0.1980 | 0.1131 | 1.9187 | 0.1019 |
| | | plastic_concentration | 1 | 0.0233 | 0.0133 | 0.4524 | 0.6444 |
| | | chlα | 1 | 0.2478 | 0.1415 | 4.8023 | 0.0070 |
| | | plastic_type:plastic_concentration | 2 | 0.0956 | 0.0546 | 0.9264 | 0.4565 |
| | Day 56 | plastic_type | 2 | 0.2483 | 0.1338 | 2.2851 | 0.0849 |
| | | plastic_concentration | 1 | 0.0543 | 0.0293 | 1.0003 | 0.3676 |
| | | chlα | 1 | 0.1998 | 0.1077 | 3.6781 | 0.0170 |
| | | plastic_type:plastic_concentration | 2 | 0.1037 | 0.0559 | 0.9539 | 0.4266 |
| | Day 94 | plastic_type | 2 | 0.1176 | 0.6539 | 1.2036 | 0.2917 |
| | | plastic_concentration | 1 | 0.1393 | 0.0775 | 2.8528 | 0.0410 |
| | | chlα | 1 | 0.1490 | 0.0829 | 3.0512 | 0.0739 |
| | | plastic_type:plastic_concentration | 2 | 0.2689 | 0.1495 | 2.7526 | 0.0320 |

*Metric* refers to the dataset analyzed (16S or 18S). *Time* indicates the specific sampling period on which the analysis was conducted. *Effect* is the explanatory variable tested (e.g., plastic type, concentration). *df* is the degrees of freedom associated with each effect. *SumofSqs* is the sum of squares attributed to the effect. $R^2$ represents the proportion of variance in community composition explained by the effect. *F* is the F-statistic from the PERMANOVA test. *p-value* indicates the significance of the effect on that date.

**Supplementary Table 8** Community metrics for no-plastic control tanks.

| *Dataset* | $R^2$ | *F* | *p* |
|---|---|---|---|
| *Zoop Abundance* | | | |
| PERMANOVA | 0.270 | 1.666 | 0.095 |
| ANOVA | | 2.511 | 0.136 |
| Shannon | | 0.816 | 0.472 |
| *16S* | | | |
| PERMANOVA | 0.240 | 1.898 | 0.053 |
| ANOVA | | 1.000 | 0.397 |
| Shannon | | 0.290 | 0.753 |
| *18S* | | | |
| PERMANOVA | 0.200 | 1.421 | 0.250 |
| ANOVA | | 0.411 | 0.673 |
| Shannon | | 0.855 | 0.452 |

Results show variation over time in control tanks based on PERMANOVA (community composition), ANOVA on total abundance and Shannon diversity. $R^2$, F-statistics, and p-values are reported for each test. These analyses assess background temporal variation in the absence of plastic exposure.

**Supplementary Table 9** Water quality summary for all ponds on day 0

| *Variable* | *Min* | *Mean* | *SD* | *Max* |
|---|---|---|---|---|
| Temperature (°C) | 12.2 | 12.443 | 0.104 | 12.6 |
| Dissolved $O_2$ (%) | 88.3 | 100.763 | 5.273 | 113.9 |
| Dissolved $O_2$ (mg/L) | 9.37 | 10.726 | 0.567 | 12.22 |
| Conductivity (μS) | 0.64 | 0.672 | 0.024 | 0.754 |
| TDS (g/L) | 0.416 | 0.437 | 0.015 | 0.49 |
| Salinity (ppt) | 0.31 | 0.329 | 0.013 | 0.37 |
| pH | 9.2 | 9.433 | 0.152 | 9.78 |
| Chlorophyll-a (μg/L) | 9.732 | 29.985 | 15.907 | 71.562 |

Min, mean, standard deviation (SD), and maximum values are shown for each variable based on measurements collected from all ponds prior to the addition of plastics.
